# Evolutionary analyses guide selection of model systems to investigate proto-oncogene function in ALK and LTK

**DOI:** 10.1101/795468

**Authors:** Zheng Wang, Alex Dornburg, Junrui Wang, Elizabeth S. Mo, Francesc Lopez-Giraldez, Jeffrey P. Townsend

**Author notes:** Z.W. and A.D. contributed equally to this work.

## Abstract

Model systems to investigate oncogene-driven cancer have played an essential role in the development of therapies for cancer. However, not all systems are appropriate for all therapeutic targets. Knowing where and when proto-oncogenes and their interactors originated in evolutionary history is key to understanding which organisms can serve as models. Here we investigate two tyrosine kinase receptors that underlie tumorigenesis in cancer: anaplastic lymphoma kinase (ALK) and leukocyte tyrosine kinase (LTK). In *Drosophila melanogaster, Caenorhabitis elegans*, and *Homo sapiens*, the discovery of putative ligands Jeb, Hen-1, and AUG has the potential to accelerate the development of novel therapeutics. However, homology of these ligands and receptors is unclear. We performed an exhaustive search for their homologs spanning the metazoan tree of life. Jeb and Hen-1 were restricted to species that diverged prior to the origin of all vertebrates. No non-vertebrate species had ligands orthologous to AUG. Instead, an ancestral receptor tyrosine kinase and AUG gene were present in early vertebrates and are still solitary in lamprey; we demonstrate that the early embryonic expression of AUG in lamprey parallels its expression in model mammal systems. The presence of ALK and LTK in jawed vertebrates is an evolutionary innovation, as is a previously unrecognized functional convergence within ALK and LTK occurring between actinopterygians and sarcopterygians. Our results provide the phylogenetic context necessary for the selection of model organisms that will provide informative investigations of the biology of these critically important tyrosine kinase receptors, enabling successful therapeutic development.

**Significance:** The anaplastic lymphoma kinase ALK can be oncogenically altered to become a driver of several malignancies, including non-small-cell lung cancer and anaplastic large-cell lymphomas. The development of therapeutics targeting this gene depend on the discovery of its interacting partner ligands in relevant model organisms. ALK is found across most major animal groups including mammals, fishes, and invertebrates. Correspondly, several candidate ligands for ALK and its duplicate LTK have been advanced by research in model species. However their homology to the human ligands and therefore their potential to guide therapeutic development is unknown. Our comparative evolutionary analysis revealed which model organisms had functional receptor-ligand pairings that are informative regarding the role of these genes in human tumorigenesis.

## Introduction

Diverse non-human models are essential to investigations of the biology of key genes that promote the origin and spread of cancer. Numerous model systems spanning the Tree of Life from yeasts to vertebrates have all demonstrated high utility to reveal pivotal insights into the biology of human cancers (1–5). However, gene function and regulation evolves, leading to divergence in the utility of model systems for investigation of a given cancer gene. Divergence in the presence and absence of components of a gene network, presence and absence of relevant domains, and even functional divergence of individual sites within genes can mislead inferences about human cancer that are based on model-organism biology. This molecular divergence is especially challenging to understand when underlying molecular divergences are obscured by convergences in phenotype. The problem of distinguishing homology from convergence has grown particularly acute within receptor tyrosine kinases (RTKs) such as anaplastic lymphoma kinase, which is known to play key roles in several malignancies, including non-small-cell lung cancer and anaplastic large-cell lymphomas (6–9).

Receptor tyrosine kinases (RTKs) are a large family of cell-surface receptors with kinase activity sensitive to specific ligand signals that are found across metazoans and that play major regulatory roles in a wide range of cellular processes (10–13). Anaplastic lymphoma kinase (ALK) and leukocyte tyrosine kinase (LTK), are two well-known RTK proto-oncogenes whose roles in oncogenesis and potential as therapeutic targets have been intensively studied (9, 14–22). To date, nearly 30 ALK oncogenic fusion partner genes have been identified in multiple types of cancers (23). Several mutation types in full-length ALK are found at higher than expected prevalences in ALK-driven cancer tissues, including ligand-independent mutations (24), ligand-dependent mutations (25), and a rare kinase-dead mutation (26). In addition to these function-altering mutations, abnormal activation of ALK can be caused by overexpression (27, 28). Justified by structural conservation of ALK between invertebrate models as diverse as fruit flies, nematodes, and humans, model organisms have been central to illuminating the biology related to these oncogenic alterations. However, translation of fundamental findings in model organisms (12, 29–31) to targeted therapeutic development in humans necessitates disentangling potentially complex patterns of divergent and convergent evolution.

Three core aspects of evolutionary history require investigation to establish the extent to which model systems provide functional parallels of humans. First is the question of ALK and LTK homology: a gene that by a simple BLAST search appears to be ALK or LTK in one model organism could in fact be the other with its very divergent regulatory apparatus and functional repertoire. Second, key domains in ALK and LTK have been gained and lost and may have undergone convergent as well as divergent evolution. ALK and LTK have a tripartite structure including an intracellular kinase domain (KD), a single transmembrane domain (TMD), and an extracellular domain (ECD) (13, 32–34). Within the ECD, a low-density lipoprotein receptor class A (LDLa) repeat and two protein tyrosine phosphatase Mu (MAM) domains are also conserved, the latter playing a role in RTK homo-dimerization, which leads to rapid activation of the kinase domains across metazoans (35). Knowing the history of gain, loss, and sequence evolution of these domains is essential to knowing functional parallels between model systems and humans. Third, three ligands of Alk have been identified: Jelly belly (Jeb) in *Drosophila melanogaster*, Hesitation behavior-1 (Hen-1) in *Caenorhabitis elegans (36–38)*, and Augmentor (FAM150 or AUG-α in *Homo sapiens*) (Guan et al 2015, Reshetnyak et al. 2015). Understanding the identity and functional interactions of these ligands with their cognate receptors has been argued to be vital to the development of novel therapeutics (39–43). However, the homology of these ligands identified in diverse organisms has not been established (44) and is essential to the establishment of the translational significance of research on each ligand in its respective model organism.

In this study, we performed an exhaustive search for homologs of vertebrate ALK, LTK, and AUG against genomes of organisms that include all major vertebrate lineages, additional chordates, hemichordates, and protostomes. Across these genomes, we identified genes homologous to those known to encode ALK ligands, providing evidence for the origins of AUG ligands as a vertebrate-specific innovation. We further reconstructed ancestral sequences of vertebrate AUGs and performed phylogenetic analysis of the evolution of vertebrate ALKs, LTKs and AUGs to reveal the history of major events in their evolution, including the gains and losses of genes and the evolution of functional domains. Using the robust gene phylogenies obtained, we further identified amino acids that likely play essential roles in the functional divergence between gene paralogs. By determining the origins and evolution of the proto-oncogenic tyrosine kinases ALK and LTK and their ligands across the history of Metazoans, our results provide the necessary foundation for effective use of major model organisms and the informed investigations of the functional roles of these genes.

## Results

### Phylogenetic distribution of *ALK*/*LTK, JEB, HEN-1* and *AUG* homologs

BLAST searches (including blastp and tblastn) revealed a diversity of sequences that are potentially homologous to sequences of ALKs and associated ligands from model and non-model organisms across the genomes available at Ensembl (45) and NCBI (46); **Table S1**). ALKs identified in nematode and fruit fly genomes exhibited sufficient conservation of sequence for homologous alignment in the GR and kinase domains. Scrupulous investigation of unannotated regions of genomes resulted in the identification of a single ALK from a previously unannotated sequence in North American lamprey genome (47, 48) as a sequence of 793 amino acids, from a 48120 bp coding region (**Table S1**). No LTK was found in the North American lamprey. Searching for a similar protein in the Japanese lamprey genome (49) via tblastn yielded a hypothetical protein of 781 amino acids that can be annotated from a 50589 bp coding region (KE993688.1: 2555306-2605577). Eighteen 188–7662 bp GT-AG spliced introns were found in the *L. camtschaticum* ALK-like protein that are conserved in size and location in comparison to ALK in *P. marinus*. No LTK was found in the Japanese lamprey genome.

In a parallel of our finding of only a single ALK homolog in the lamprey genomes, only a single AUG homolog was found. No orthologs of Jeb or Hen-1 were found in any vertebrates, with Hen-1 restricted entirely to the nematodes *C. elegans* and *Loa*. In contrast, Jeb-1 was widespread among protostomes, with our identification of Jeb-like proteins in *Aplysia* extending the presence of this ligand to mollusca. As lampreys are members of the earliest-diverging lineage of living vertebrates, this finding of a single ALK and AUG suggests that a duplication of this ligand-receptor pair gave rise to LTK and a second AUG ligand after the divergence of the lamprey lineage from the vertebrate ancestor and prior to the diversification of jawed vertebrates.

The AUG-like protein had been annotated as a 146 amino-acid hypothetical protein (JL3715) in *L. camtschaticum*. Searching for a similar protein in the North American lamprey genome via tblastn resulted in a 147-amino-acid protein within a 7000-bp coding region (GL476337dna: 19000:26000). Two possible GT-AG spliced introns, spanning 2671 and 2977 bp, were also conserved in position and size in comparison to the AUG-like protein annotated in the Japanese lamprey genome. In AUG, a potential signal peptide of 23 amino acids was found to be conserved between the two lamprey genomes (**Table S1**). To verify expression of ALK and AUG transcripts in humans, we used PCR-based (RACE) strategies in muscle, brain, liver, and eye tissues from adult and ammocoete lampreys (Table S2) combined with analysis of RNA sequencing data from eight transcriptomic projects of *P. marinus* in NCBI Sequence Read Archive (SRA) that mapped to the genomic coding region for the annotated AUG (**Table S3**). 11641 reads from 53 samples that were collected from embryos 2.5–5 days post-fertilization that perfectly matched expected transcripts from our annotation of AUG in *P. marinus*. Tissue-specific expression patterns of AUG largely matched expression of ALK. AUG transcripts were found in the neurula stage, olfactory tissues after exposure to copper, meiotic testes and brains, and samples with post-injury spinal cord and brain tissue; ALK transcripts were detected in 83 samples, including all tissues where AUG was detected, plus embryonic tissues sampled ≤5 days post-fertilization and brain, liver, and kidney tissues of later developmental stages (both larval and adults). Expression of AUG was not detected from liver tissues from lamprey specimens at both the ammocoete and adult life stage.

Our inferred phylogenetic relationships of these sequences reflects our findings of homology and challenge current conceptions of ligand-receptor homology (**Fig. 1**). Orthologs of ALK are present in almost all vertebrate genomes as well as most non-vertebrate groups. However, ALK appears to have been repeatedly lost in early diverging chordate lineages, including *Ciona, Branchiostoma*, and *Eptatretus* (**Fig. 1**). LTK presumably arose from a duplication at the common ancestor of *Petromyzon* and jawed vertebrates (48, 50)—simultaneous with the appearance of both AUG ligands (**Fig. 1**). LTK appears to have subsequently been lost independently in monotremes and tenrecs (**Fig. 1**). Our identification of Jeb in *Strongylocentrotus* marks the first identification of this protein in a deuterostome. However, we did not find a Jeb homolog in any chordate lineage, indicating that Jeb was lost prior to the diversification of vertebrates (**Fig. 1**). Our phylogenetic analyses provide strong evidence that among the candidate ligands of ALK and LTK receptors, vertebrates possess only AUGs.

**Figure 1.**
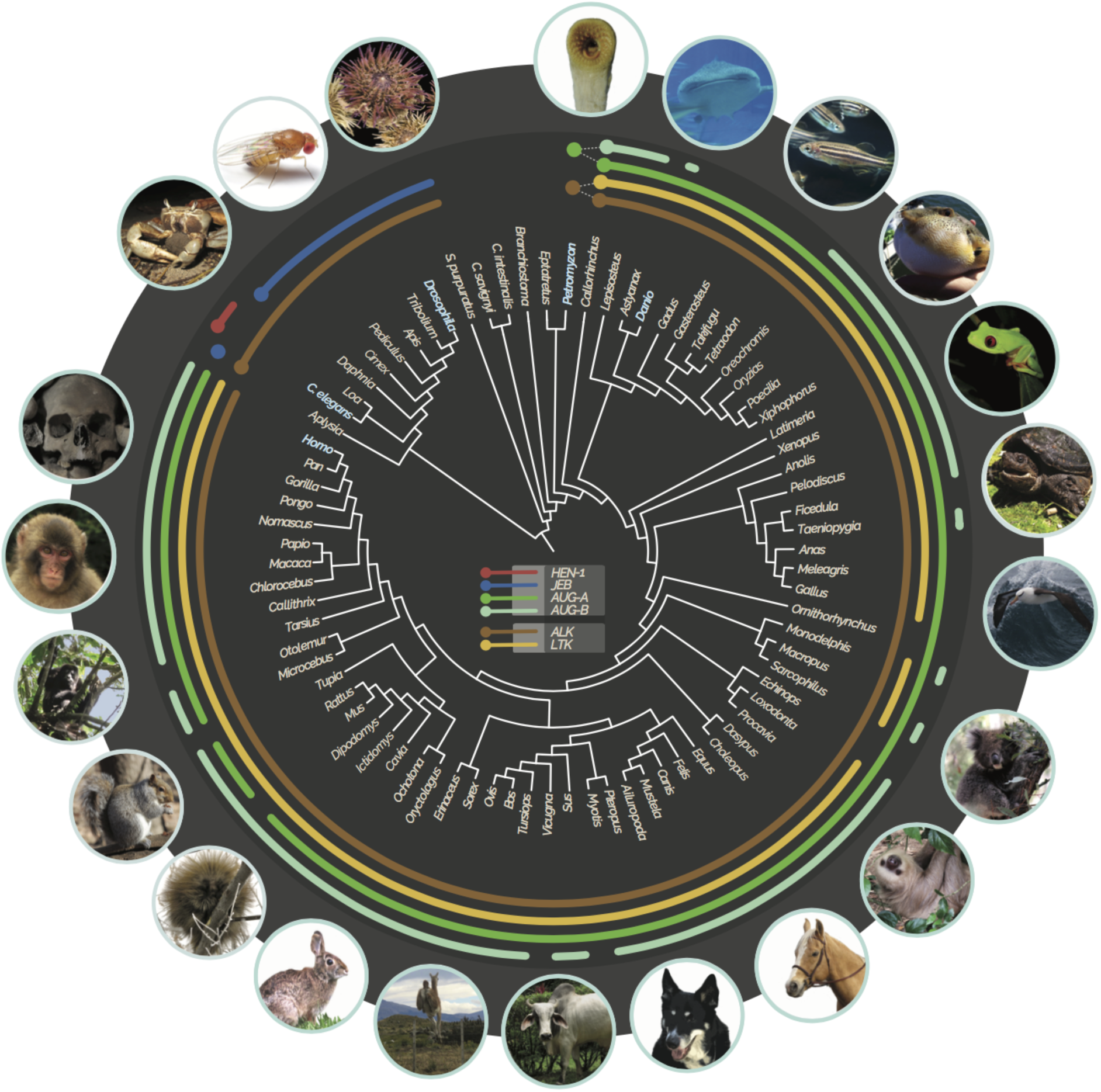
Phylogenetic distribution of HEN-1 (red), JEB (blue), ALK (brown), LTK (yellow), AUG-α (green), and AUG-β (teal) orthologs in metazoan genomes (arcs span taxa with each gene; colored taxon names correspond to model organisms discussed in the text), especially those of *C. elegans, Drosophila, Petromyzon, Danio*, and *Homo* (taxon names colored). Identification of homologous genes was confirmed with phylogenetic analyses, and gene accession numbers are provided (**Table S1;** Photo credits Dan Warren [koala], Katerina Zapfe [squirrel], Bronwyn Williams [urchin], Matt Bertone [*Drosophila*], Lynn Ketchum [zebrafish: creative commons cropped from original], and Alex Dornburg [all others]).

### The diversification of ALK and LTK in vertebrates

Phylogenetic analysis provided strong support [Bayesian posterior probability (BPP) = 1.0] for a history in which the duplication of ALK gave rise to LTK near the common ancestor of jawed vertebrates and lampreys (**Fig. 2, Fig. S1, Table S1**). Strong support of the reciprocal monophyly of jawed vertebrate ALK and LTK across jawed vertebrates (BPP = 1.00) enables comparisons of the evolutionary history of ALK & LTK that demonstrate divergent patterns of domain acquisitions and losses, as well as divergent rates of molecular evolution for these genes (**Fig. 2**). For example, several non-mammals exhibit a signature of accelerated evolution of nonsynonymous substitution in ALK, an acceleration that contrasts with a significantly faster rate of molecular divergence of mammal LTKs compared with ALK (**Fig. 2**; **Tables S4 and S5**). We found that both ALK (Iss = 1.08, Iss.c = 0.83, *P* < 0.01; **Fig. S5**) and LTK (Iss = 1.21, Iss.c = 0.83, *P* < 0.01; **Fig. S5**) have experienced severe substitution saturation (51, DAMBE; 52) and exhibit a sharp decline of phylogenetic informativeness (PhyInformR; 53) even within the divergence of mammals. Rates of evolution of ALK and LTK are contrasting between “fish” (ray-finned fishes, sharks, and Coelacanth) and mammals: mammal LTK exhibits significant saturation (Iss = 1.07, Iss.c = 0.84, *P* < 0.01) relative to “fish” (Iss = 0.67, Iss.c = 0.84); and ALK for “fish” exhibits significant saturation (Iss = 1.07, Iss.c = 0.84, *P* < 0.01) relative to mammals (Iss = 0.65, Iss.c = 0.84).

**Figure 2.**
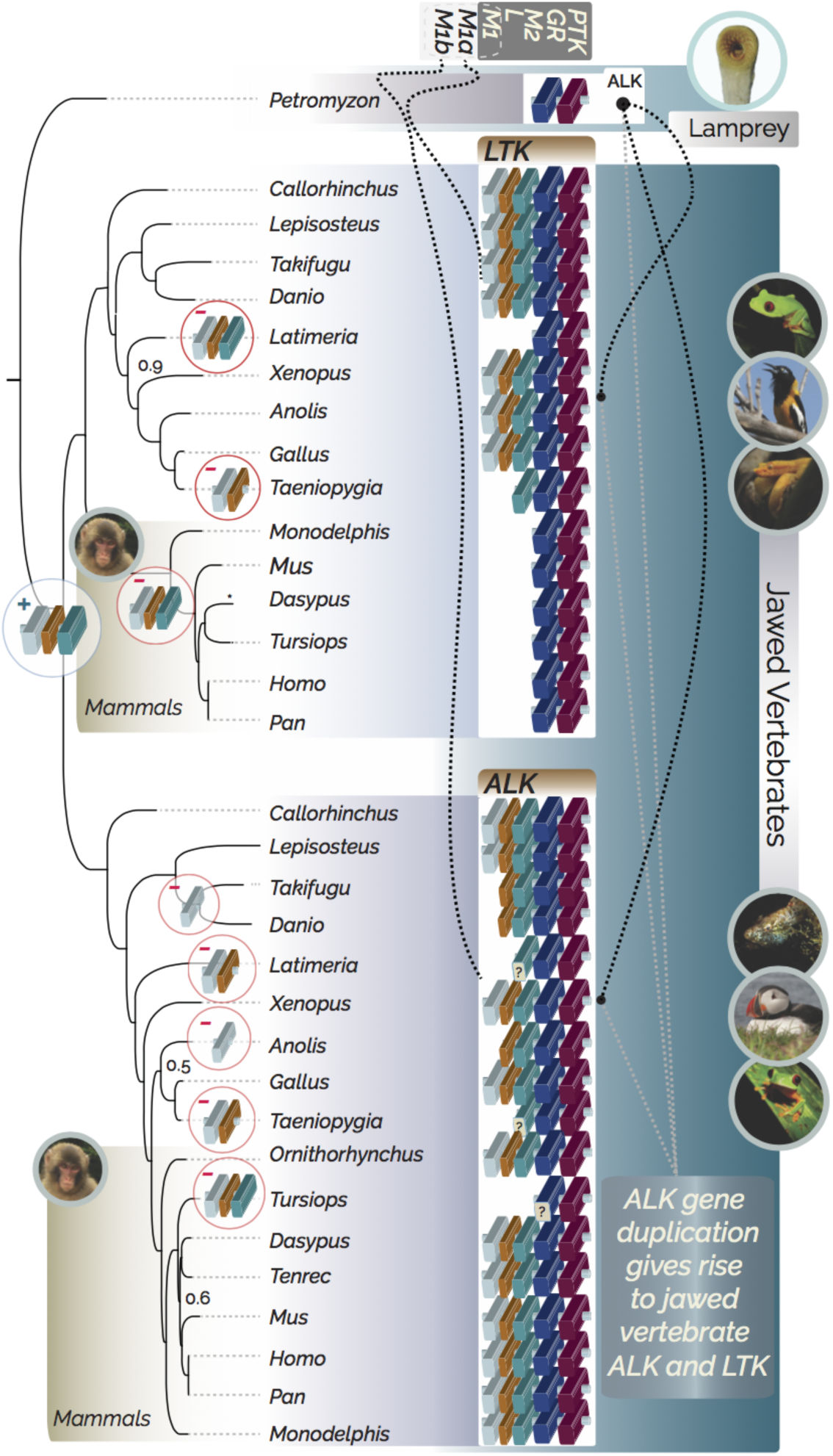
Phylogeny of vertebrate ALKs and LTKs. A simple ALK in Lamprey (*Petromyzon*) features only PTK and GR domains (blue and red blocks). M_2_, L, and possibly M_1_ domains (teal, orange, and gray blocks) are depicted as a gain (+) that was duplicated in jawed vertebrates, giving rise to ALK and LTK (black dashed lines from simple ALK, indicated with gray lines from inset text). Dashed lines from MAM domains indicate the presence of M_1a_ and M_1b_ in ALK and LTK, respectively; the M_1a_ and M_1b_ domains exhibit no discernable sequence similarity (“?” on teal block at the base of jawed vertebrates). Some domains in some lineages exhibited low levels of sequence similarity to domains observed in other vertebrates (“?”s on GR block on the lineage to *Tursiops*, and M2 on the lineage to both *Latimeria* and *Taeniopygia*). Domain losses (-) are indicated at internodes where they were reconstructed to occur (e.g. loss of M1, L, and M2 in LTK in mammals—highlighted with a gold gradient, and loss of M1, L, and M2 in *Tursiops* in ALK). Unlabeled internodes all exhibited strong statistical support (Bayesian Posterior Probability [BPP] > 0.98). Labeled internodes indicate level of support for the most probable topology. Photos: AD.

Placement of the evolution of ALK and LTK into a phylogenetic context further enables evaluation of the homology of structural domains observed within these proteins. For example, MAM domains, which are present in fruit fly ALK, are not homologous to those found in ALK or LTK in humans. For vertebrate ALKs, there is no sequence similarity between MAM1 and MAM2 domains in vertebrate ALKs or and between MAM1 in ALKs and MAM1 in LTKs. Ancestral reconstruction of MAM domains in vertebrate ALK and LTK using `BayesTraits` 1.0 (54) strongly (BPP > 0.95) supported the origination of the MAM2 domains in the most recent common ancestor of vertebrates, and their subsequent loss in mammal LTKs (**Fig. 2, Fig. S2**). Similarly, MAM1 domains are likely the product of parallel evolution within jawed vertebrate ALK and LTK.

We found evidence for functional divergence in different sites between ALK and LTK, with six sites (745-Lys, 760-Leu, 767-Lys, 795-Ile, 808-Asn, and 863-Asn positions of human ALK) identified to be of significant (*P* < 0.05) importance in the differential function of ALK and LTK (**Fig. S3**). These sites were all located between the MAM2 domain and the GR region, with the exception of 863-Asn, located in the GlyR domain. Functional divergence analyses further predicted 11 amino acids in human LTK to be of significant (*P* < 0.01) importance in the differential function of LTK between mammals and non-mammals groups (98-Thr, 120-Leu, 152-Leu, 171-Gly, 200-Gly located before the GlyR domain and 216-Tyr, 226-Glu, 245-Arg, 261-Ala, 262-Pro, 267-Arg located within the GlyR domain), five of which are conserved across ALK and LTK in jawed vertebrates (**Fig. S4**).

### The evolution of augmentor (AUG) in vertebrates

Investigation of transcriptomes and sequenced genomes revealed that AUG is an innovation shared by all vertebrates, potentially synchronous in origin with the hypothesized early vertebrate genome duplication that may have given rise to ALK. Using the annotated lamprey AUG sequence and the maximum likelihood ancestral sequence for the most recent common ancestor of vertebrate AUG and lamprey AUG revealed no potential homologs in searches of any non-vertebrate deuterostome, protostome, or non Metazoan genomes. Only two sequences, present within a 40-amino-acid stretch, produced a significant alignment to sequences from two aquatic cyanobacteria, *Gloeobacter violaceus* and *Oscillatoria acuminata*—speculatively implicating an ancient horizontal gene transfer between these aquatic cyanobacteria and lamprey. Our reconstruction of the evolutionary history of AUG strongly supports its duplication into AUG*-α* and AUG*-β* prior to the most recent common ancestor of jawed vertebrates (**Figs. 1, 3**). These two lineages have since been heterogeneously maintained across vertebrates (**Fig. 1)**, and differ from lamprey AUG in structure; lamprey AUG only encodes three cysteines near the C-terminus (**Fig. 3C**). Functional divergence analysis further identified three sites in human AUG*-α* (81-Glu, 91-Leu, and 146-Val) as being potentially important to the differential function of the AUG paralogs.

**Figure 3.**
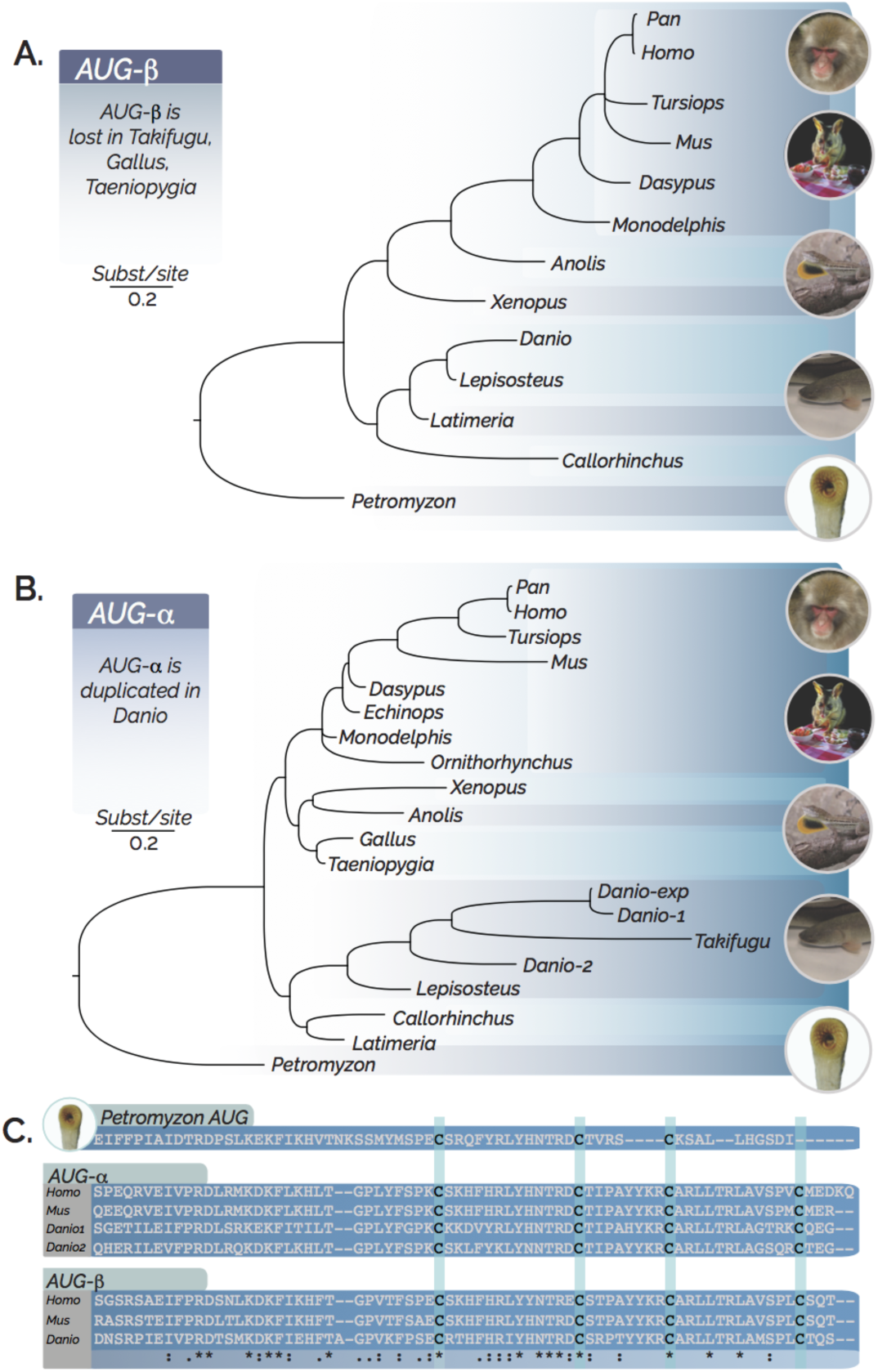
Phylogeny of vertebrate A) AUG-β and B) AUG-α, demonstrating faster-evolving AUG-α in lineages with long branches, illustrated with C) an alignment of selected AUGs demonstrating sequence conservation between lamprey AUG and its homologs in mammals and zebrafish. Key cysteine positions near the C-terminus are highlighted in green. Labels at branches indicate Bayesian posterior probabilities < 0.98. Images: AD (Possum courtesy of Dan Warren).

Orthologs of AUG*-α* exhibit accelerated evolution (high non-synonymous substitution) across jawed vertebrates, suggesting a comparatively conserved function of AUG*-β* (**Table 1**). This hypothesis is supported by a relative ratio test demonstrating significantly greater numbers of amino acid substitutions in AUG-α than in AUG-β (*P* < 0.05). Both AUG-α (Iss = 1.39; Iss.c = 0.76) and AUG-β (Iss = 1.45; Iss.c = 0.76) have experienced severe substitution saturation (DAMBE; 51, 52) and exhibit a sharp decline of phylogenetic informativeness (PhyInformR; 53) at timescales corresponding with the divergence of mammals (**Fig. S5**).

## Discussion

Here we have revealed that the augmentor ligands are an innovation shared by all vertebrates. Our results place the origin of AUG synchronous with a hypothesized early vertebrate genome duplication and demonstrates that JEB and HEN-1 are not homologs of AUG. We showed that after the genesis of AUG, an additional duplication of AUG likely occurred in the most recent common ancestor of jawed vertebrates, giving rise to the ligands AUGα and AUGβ. This duplication of AUG coincided with an additional duplication of anaplastic lymphoma kinase (ALK) forming the leukocyte tyrosine kinases (LTK). Ancestral state reconstructions of vertebrate ALKs and LTKs revealed both clade- and gene-specific gains and losses of MAM domains, supporting functional convergence between non-tetrapod LTK (such as in zebrafish) and mammal ALK (such as in humans). Consideration of the complexity of subsequent evolution of ALK and LTK lineages reveals the utility of the lamprey as the simplest model system for study of ALK and AUG receptor-ligand biochemistry and molecular genetics, and facilitates the interpretation of the molecular biology of these genes in vertebrate models of human cancer.

The deep evolutionary split between ALK-bearing vertebrates and ALK-bearing protostomes corresponds not only to a divergence in receptor-ligand association, but also to a greater functional divergence of ALK between the two lineages that renders research progress on multiple fronts with diverse model systems challenging. For instance, ALK plays a central role in the formation of the visceral gut formation, growth, and neurogenesis in protostomes (55, 56), and a role in neuronal proliferation, differentiation, and survival in vertebrates (34, 57). The lineage-specific evolution of ALK function across Metazoa likely underlies the diversity of hypotheses regarding how human functions and phenotypes of ALK homologues relate to the functions and phenotypes observed in protostome models such as nematodes and fruit flies. These divergent functions and phenotypes include the binding specificity between the receptor and ligand, the suppression of the receptor, and the function of domain structures (36, 58, 59). Functional roles of genes often shift when genes duplicate, necessitating evolutionary analysis with inclusive data sets (60–63). In the case of ALK, as in other mysteries of vertebrate molecular evolution, the jawless lamprey genome held essential clues to the origins and evolution of genes, gene families, and other genetic elements in jawed vertebrates (64, 65). Accordingly, some of the functional shift between protostome ALK and vertebrate ALK can be attributed to the consequences of ancient duplication of ALK into ALK and LTK following the early vertebrate genome duplication. Our findings represent a prime example of how the lamprey genomes can reveal the origins of vertebrate-specific traits (66–68). The early evolutionary divergence of lampreys from the majority of living vertebrates (69–72) has repeatedly illuminated highly conserved elements of vertebrate molecular biology including basic developmental processes such as how the brain and nerves develop and revealing genes whose activity aids spinal cord healing (73).

Structurally, we found lamprey ALK is similar to mammal LTK: both feature the PTK and GR domains and lack MAM and LDL domains. Our analysis of transcriptomic data with newly annotated lamprey ALK and AUG demonstrated that ALK and AUG are coexpressed post-fertilization, during early embryonic development or after injury to nerve or brain tissues, which suggests a role of ALK and AUG in nerve and brain development. Expression of ALK in mice and humans is also highest during embryonic development, quickly drops after birth, and is maintained at a low level (74, 75), suggesting the functional roles of ALK and AUG to be conserved across vertebrates. Biophysical binding data suggests AUG-α is a dual-specific ligand for both ALK and LTK while (29) AUG-β is a monospecific ligand of only LTK (29). However, the physiological functions and interactions of these two ligands with receptors appear highly heterogeneous between species (e.g. 12, 43, 58, 76) For instance, although the expression patterns of ALK receptors and AUG ligands between lamprey and mammals are similar, lamprey AUG is structurally different. Lamprey AUG has only three C-terminal cysteines, instead of the four cysteines found in other vertebrates. Of those three C-terminal cysteines, the first two are in locations conserved across all other vertebrates. Cysteines form inter- and intra-molecular disulfide bonds that are important to ligand structural integrity (12, 77), suggesting that lamprey provides a key reference species for further study of the functional roles of the third and fourth cysteines of vertebrate AUGs.

The duplication of ALK into ALK and LTK following the divergence of lamprey with the most common recent common ancestor of jawed vertebrates constitutes a vertebrate-specific evolutionary novelty that, despite ancient divergences, has maintained striking functional similarities between these two lineages of tyrosine kinases. There are numerous functional similarities within ALK, within LTK, and between ALK and LTK encourages use of a wide range of candidate vertebrate model species for investigation of the receptors, ligands, and their interactions. For instance, the LTKs of non-tetrapods, including models such as zebrafish, exhibit a strong signature of shared domain structure with mammal ALK. The only notable difference is the sequence divergence between the N-termini of ALK and LTK within the first MAM domain. Based on our ancestral state reconstructions of the LTK and ALK MAM domains, it is likely that this large sequence divergence is a consequence of convergent gains of the first MAM domain in non-tetrapod LTK and mammal ALK. In addition to these shared domains, we also found evidence for the conservation of 11 key amino acids between non-tetrapod and mammal LTK. The amino-acid identity of five of these eleven amino acids is—remarkably—also shared between non-tetrapod LTK and mammal ALK. This conservation is encouraging and consistent with previous research emphasizing the importance of the nonhuman models in ALK tumorigenesis (19). Experimental research investigating the effects of induced point mutations in the ALK and LTK coding sequences at the sites identified here would be especially likely to reveal functionally divergent aspects of ALK and LTK signaling among humans and relevant model species.

Well-developed phylogenetic and evolutionary biology application provide robust approaches to analyze large data sets for evolution of gene and gene family (78–80). Our phylogenetic analyses of AUG-α revealed an accelerated evolutionary rate that is unexpected for proteins executing critical biological functions. Lower rates of sequence evolution are typically expected for proteins believed to have a collocalized dual specificity of interaction between genes, as is the case for AUG-α interacting with ALK and LTK, based on biophysical experiments on mouse cell lines (29). Our results demonstrate that this dual specificity has not constrained the evolution of AUG-α to a slower substitution rate within mammals. Instead, molecular rates of AUG-α exceed those estimated for AUG-β, which is monospecific for LTK (29). The high substitution rates observed in mammal LTK, non-tetrapod ALK, and jawed vertebrate AUG-α could indicate increased functional specificity and lower promiscuity of interaction in these genes. In contrast, we found AUG-β homologs to be more conserved than AUG-α homologs, a signature consistent with expectations of co-evolution between signaling and receiving molecules (81–83).

The comparatively lower substitution rate in mammal ALK is consistent with evolutionary expectations of an “orphan receptor”. In mammals, ALK is activated via both ligand-dependent (29, 84) and ligand-independent (85, 86) processes, implying multiple functions of ALK and high interaction specificity between ALK and its ligand(s). These lower substitution rates potentially indicate promiscuous interactions among AUG-β and its receptors. Biological relevance of these interactions has been indicated by research on both zebrafish development and human cancers (12, 29, 44).

The differences in evolutionary rates between AUG-α & AUG-β, mammal LTK & non-tetrapod ALK, and mammal ALK & non-tetrapod LTK represent evolutionary trade-offs between functional specificity and the number of interactors a protein can achieve (87). Rapidly evolving proteins have been shown to exhibit greater functional specificity—for example, higher tissue specificity or higher promoter methylation in mammals (87). If the different rates of evolution of these receptors and their ligands are the outcomes of evolutionary trade-offs, we might expect a higher complexity of the protein networks associated with mammal LTK and non-tetrapod ALK, and expect more functional generality in networks associated with mammal ALK and non-tetrapod LTK. Further molecular biological research is warranted investigating consequent hypotheses: whether mammal LTK and non-tetrapod ALK regulate nerve development in conjunction with many highly specialized partner proteins, and whether mammal ALK and non-tetrapod LTK play broadly important and general roles in internal developmental signaling. Additional collection of data on genome-wide or gene-specific spatial and temporal co-expression in vertebrates could provide additional insight into regulatory gene interaction networks, providing hypotheses for protein-protein interaction experiments and revealing evolutionary change in the structure and function of the ALK, LTK, and AUG signaling networks. Further analysis of the function of these genes within their protein interaction networks, cognizant of their evolutionary history, could illuminate how they are co-opted in tumorigenesis and cancer progression. Unveiling the evolutionary history of these ligands and receptors will guide informative research toward an understanding of the cellular function of ALK and LTK in human cancers.

## Materials and methods

### Identification homologs of ALK and its possible ligand in non-models

To investigate homologs of vertebrate ALKs, LTKs, and their associated ligands across model organisms spanning the protostomes and deuterostomes, BLASTp and tBLASTn searches of protein sequences derived from the genomes of zebrafish (*Danio rerio*), fruit fly (*Drosophila melanogaster*), and nematode (*Caenorhabditis elegans*) were conducted against available genomes from genome databases in NCBI (www.ncbi.nlm.nih.gov) and Ensembl (www.ensembl.org; **Table S1**). Sequences with the mutually best matches between two sequences in genome pairs via BLAST search (88), were subjected to further phylogenetic analyses to confirm their homology with annotated ALK, LTK or ligand proteins. To investigate the evolution of ALK/LTK and associated AUG in vertebrates, sequences from model species that included zebrafish, frog (*Xenopus tropicalis*), chicken (*Gallus gallus*), zebra finch (*Taeniopygia guttata*), mouse (*Mus musculus*) and human (*Homo sapiens*) were queried against non-model vertebrate genomes (**Table S1**) using best-hit reciprocal BLAST searches. To illuminate the origin of vertebrate ALK/LTK and AUG, special attention was devoted to thoroughly ascertain the presence of ALK and possible ligand(s) in the genomes of the jawless vertebrates (hagfish and lamprey), as these animals represent the earliest-diverging extant vertebrate lineage (89). No sequence in the hagfish genome exhibited any similarity to vertebrate ALK, LTK, or AUG. In contrast, LTK had been identified and annotated in the Atlantic lamprey genome. Additionally, an unidentified protein annotated as hypothetical exhibited a high similarity to partial zebrafish AUG in the Japanese lamprey genome. BLAST searches of multiple ALK and AUG sequences from non-tetrapod genomes recovered highly conserved regions between the two lamprey genomes indicating the presence of homologs of jawed-vertebrate ALK and AUG.

Searches for ALK, LTK and paired ligands were additionally conducted against transcriptomes of sea lampreys (*Petromyzon marinus*), including 86 publicly available transcriptomes (**Table S3**). RNAs were additionally sampled from tissues of an ammocoete and an adult sea lamprey. The ammocoete lamprey was flash-frozen in liquid nitrogen before tissues of head, muscle, and viscera were dissected for RNA extraction. Tissues of the large adult lamprey were resected from muscle, eyes, liver, brain, and heart. All tissues were preserved in RNA*later*, then maintained at −76 C prior to RNA extraction. Total RNA was extracted from homogenized tissue with TRI REAGENT (Molecular Research Center). Messenger RNA was purified using Dynabeads oligo(dT) magnetic separation (Invitrogen). A cDNA library was generated using a SMARTer 5’/3’ RACE Kit (Takara cat no. 634860) as per the manufacturer’s instructions. First-Strand cDNA synthesis was performed using 11 uL of RNA extract and 1 uL of 3’-CDS Primer A. Rapid amplification of cDNA ends (RACE) was also performed as per the manufacturer’s instructions with custom gene-specific primers (GSPs, **Table S2**).

The genome sequence of the lamprey *P. marinus* was downloaded from Ensembl (90), and used as the reference sequence for `HISAT2` (91) to build the index and perform read alignment. Transcripts were assembled and gene-expression levels were quantified using `StringTie` (92). Sequence read data totaled 323 Gb, and the largest single dataset (based on paired-end sequencing with the Illumina HiSeq 4000 platform) amounted to 21.7 Gb. We extracted `StringTie` annotation results without a reference, based on the scaffold GL476336:19708–25867 where AUG is predicted. We manually annotated lamprey AUG and ALK with flanking regions (AUG, GL476337:10–26043; ALK, GL476342:132396–213117) and used these manually annotated sequences as references for read mapping, specifying the HHEX (93) gene as a control. Due to its high efficiency, `HISAT2` (91) was chosen to map the reads, which were subsequently extracted with `SAMtools` (94) with parameter setting `-F 4` that filters unmapped reads. Across all datasets (**Table S3**), a total of 225,182 reads mapped to sea lamprey AUG coding and non-coding sequence (26033 bp, 138× coverage), of which 11,641 reads mapped to AUG exons (441 bp, 422×). These reads were derived from samples of whole embryos 1–2.5 days post-fertilization; neural crest, kidney, brain, liver and olfactory tissue after 24 h exposure to 5 g/mL, 10 g/mL, and 30 g/mL of copper; and brains (whole brains without the olfactory lobes) and spinal cords (1 cm surrounding the lesion), harvested from 6 h to 12 weeks post injury of lamprey specimens (**Table S3**). The 11641 reads assembled into a contig that perfectly matched the annotated AUG. Reads were found at high coverage (221×–1023×) for the first 3 exons. Only one solitary read—in the SRR2238808 run (95)—mapped to the fourth exon, from the sequencing of early embryo stages 23–25. We detected eight isolated contigs in non-exon regions using `Sequencher` 5.4.6 (96) with default settings. Within these putative non-exonic regions, we found portions exhibiting high sequence similarity with homeobox proteins (HOX) and the neurotrophin type 2 gene (suggesting high potential for further gene discovery in lamprey genomes; **Table S1**). We observed no alternative splicing of AUG exons in these lamprey transcriptomes. For ALK, 6,936,551 reads were found to map onto the region spanning from the flanking region before the first exon to the flanking region after the last exon (80721 bases, 1375×), of which 1,311,610 reads mapped to the identified ALK exons (2379 bases, 8821×), representing reads that were mostly from the same samples that AUG expression was detected in: early embryo development, copper treatments, and nerve injury (**Table S3**). We detected five isolated contigs in non-exon regions. Portions of these five contigs exhibited high sequence similarity with homeobox proteins (HOX), myosin, or lamprin coding regions, again indicating significant potential for undescribed genes in lampreys (**Table S1**). Two mRNAs were observed, one that spliced out intron #18, and one that did not. Alternative splicing is known for human LTK (97).

### Molecular Phylogeny

We obtained amino acid sequences and nucleotide sequences of *ALK* / *LTK* and *AUG-α* / *AUG-β* genes from NCBI and Ensemble respectively (**Table S1**). The amino acid sequences of ALK and Jeb in *Drosophila melanogaster*, and SCD-2 (homologous with ALK in *Drosophila*) and Hen-1 in *Caenorhabditis elegans* were accessed for sequence comparison to lamprey ALK and AUG. We subsequently used lamprey sequences as outgroups for phylogenetic inference of ALK/LTK and AUG evolution in jawed vertebrates. Amino acid sequences were aligned using `MAFFT` in `Saté-II` (98, 99), while nucleotide sequences were aligned based on amino acid sequences by TranslatorX server online (http://translatorx.co.uk/). We inferred phylogenies of ALK and AUG using Markov chain Monte Carlo (MCMC) methods implemented in `MrBayes` 3.2 (Ronquist et al. 2011). Our Bayesian phylogenetic analyses were executed for 10,000,000 generations, sampling every 1,000 generations with four chains. We set lamprey ALK and AUG as outgroups for each analysis and discarded 2500 (25%) of the 10,000 trees as burn-in. We assessed convergence of the chains by quantifying potential scale reduction factors (PSF = 1.0). We visually compared computed log likelihoods across chains to confirm stationarity. For nucleotide sequences, the GTR + I + G substitution model was specified. For amino acid sequences, a mixed model of amino acid substitution was specified that allowed each amino acid model to contribute in relation to its posterior probability. We deemed branches that exhibited a posterior probability (PP) higher than 0.98 to be strongly supported (**Figs. 2**–**3, Fig. S1**). We conducted analyses on both the aligned amino-acid and nucleotide sequences separately.

### Ancestral reconstruction

Using our phylogeny, ancestral sequence reconstructions were performed using both likelihood and Bayesian approaches as implemented in `PAML` 4 (100). We used `codeml` to conduct codon-based ancestral sequence reconstruction of the common ancestor of mammals as well as all vertebrates. Reconstructed sequences of ALK and AUG for the common ancestor of lamprey and jawed vertebrates were then used to search against non-vertebrate genomes for possible homologs. Ancestral states of MAM domains in vertebrate genomes were estimated using maximum likelihood (ML) criteria in `BayesTraits` (101, 102). We coded the presence or absence of MAM domains for ALK and LTK, and used the multiState method of discrete character evolution to reconstruct gains or losses of MAM domains (**Fig. S2**).

### Selection tests

To test for positive selection along specific branches, we used branch models implemented in `PAML` (100, 103), in which the ratio of nonsynonymous to synonymous substitution (*ω*) was allowed to vary among branches in the phylogeny (**Table S4, S5**). The ratio *ω* was estimated for the branch of interest (the ‘foreground’ branch) and the rest of the tree (the ‘background’) in the phylogeny reconstructed from a multiple sequence alignment. To evaluate whether there was a statistically significant difference between the branch model and the null model, a likelihood ratio test (LRT) was applied. To search for positively selected sites, site models permitting *ω* to vary among sites were used. We set NSsites to equal 0, 1, 2, 7, and 8, then conducted likelihood ratio tests between pairs of the models to identify the best fitting model comparing M_1a_ (Nearly Neutral) against M_2a_ (Positive Selection), and M_7_ (beta) against M_8_ (beta & *ω*), each with two degrees of freedom. A Bayes-Empirical Bayes analysis was performed to identify the sites evolving under significant positive selection (104). Clade model C was fit to the data to evaluate whether *ω* differed between clades ALK and LTK and between clades AUG*-α* and AUG*-β* in mammals (104, 105). The improved model M_2a_rel_ was used as the null model for the likelihood ratio test on clade model C results (106).

The degree of saturation of substitutions for ALK, LTK, and AUG proteins was assessed by `DAMBE` (51) and by inspection of phylogenetic informativeness profiles (107) visualized with the `R` package `PhyInformR` (53). To estimate site rates, we used `HyPhy` (108) in the `PhyDesign` web interface (109). PI profiles were depicted along a relative ultrametric guide topology generated in `BEAST` v. 2.4.7 (110) with a prior root height of 1.0. At the depths of divergence examined exhibited some evidence of saturation with regard to substitutions, we compared their maximum likelihood rates of sequence evolution using a Likelihood Ratio Test conducted in `Hyphy` (108), enabling meaningful comparisons of relative rate differences between AUG paralogs.

### Functional divergence analysis

We used `DIVERGE` 3.0 (111) to test for functional divergence of the gene pairs. `DIVERGE` tests for site-specific shifts in evolutionary rates after gene duplication or speciation. The coefficient of divergence (*θ*_*D*_) was calculated to test against a null hypothesis of no functional divergence between ALK and LTK, between mammal ALK and fish ALK, between mammal LTK and fish LTK, and between AUG*-α* and AUG*-β*. We employed the default posterior probability cutoff of 0.5 for detection of site-specific shifted evolutionary rates (111). Amino acids with significant (*P* < 0.05) roles in functional divergence between gene paralogs were predicted.

## Acknowledgments

We thank G. J. Watkins-Colwell at the Peabody Museum of Natural History for help collecting lamprey specimens and tissue samples, and J. Yoder, K. Zapf, and L. Abrams for valuable discussions and critical comments. This research was supported by National Science Foundation IOS-1755242 to AD and by research support from the Notsew Orm Sands Foundation to JPT.

