## supplemental data in one file for "Evolutionary analyses guide selection of model systems to investigate proto-oncogene function in ALK and LTK" (from another version of this preprint: version 1, 2019-10-07)

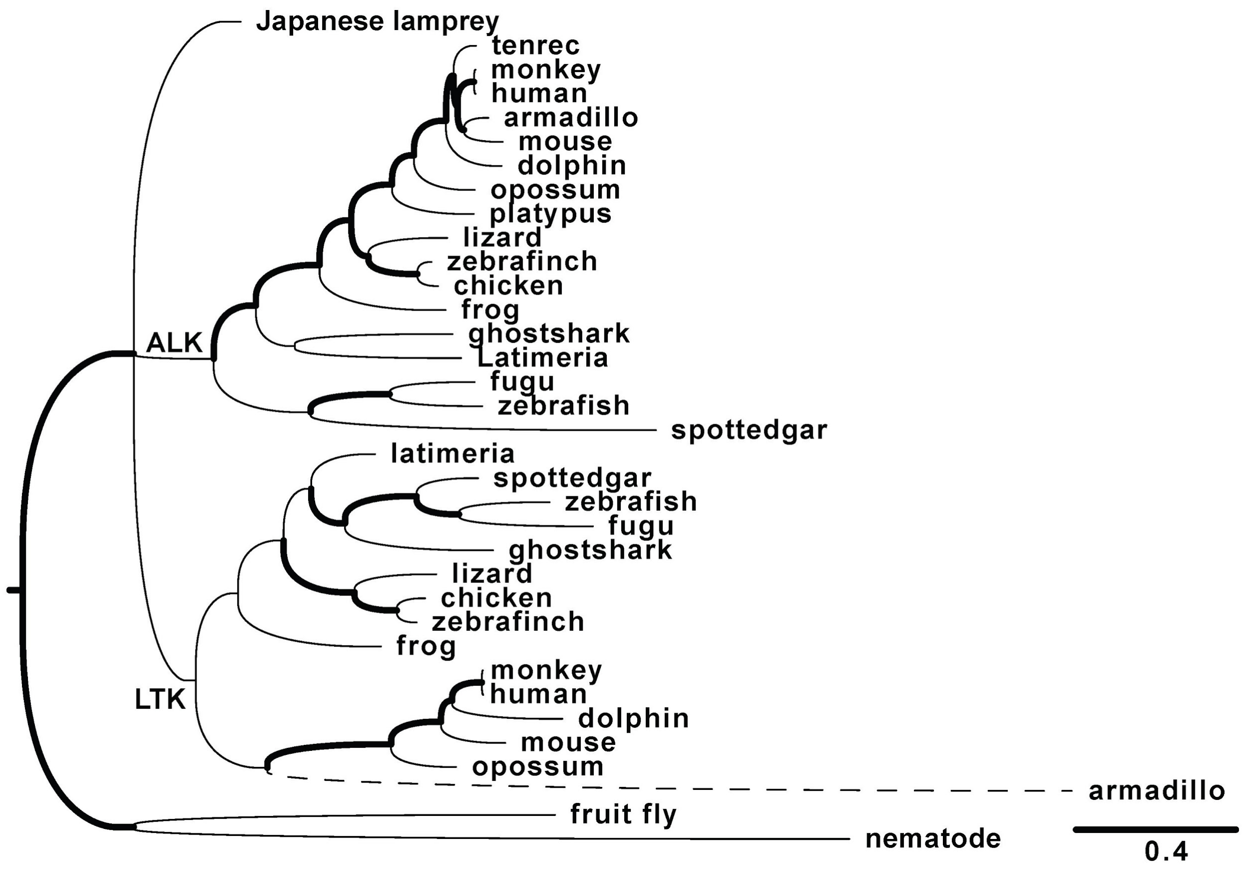


**Figure S1.** Phylogeny of vertebrate ALK and LTK, rooted with fruit fly *ALK* and the nematode homolog *SCD2*. The phylogeny was inferred from a joint alignment of ALK and LTK using `MrBayes` (112, 113). Branches with strong support (PP > 0.98) are in boldface. An extremely long branch extending to armadillo LTK (dashed line) was shortened for clarity of display of the tree topology and the lengths of other branches.

| | M1 Absence | M1b | M1a |
|---|---|---|---|
| Node A | 0.85 | 0.12 | 0.03 |
| Node B | 0.001345 | 3.40E-05 | 0.999 |
| Node C | 0.108 | 0.889 | 0.0024 |
| Node D | 0.26 | 0.19 | 0.55 |
| Root | 0.51 | 0.22 | 0.27 |

| | M2 Absence | M2 |
|---|---|---|
| Node A | 0.99 | 0.01 |
| Node B | 0.01 | 0.99 |
| Node C | 0.01 | 0.99 |
| Node D | 0.05 | 0.95 |
| Root | 0.60 | 0.40 |

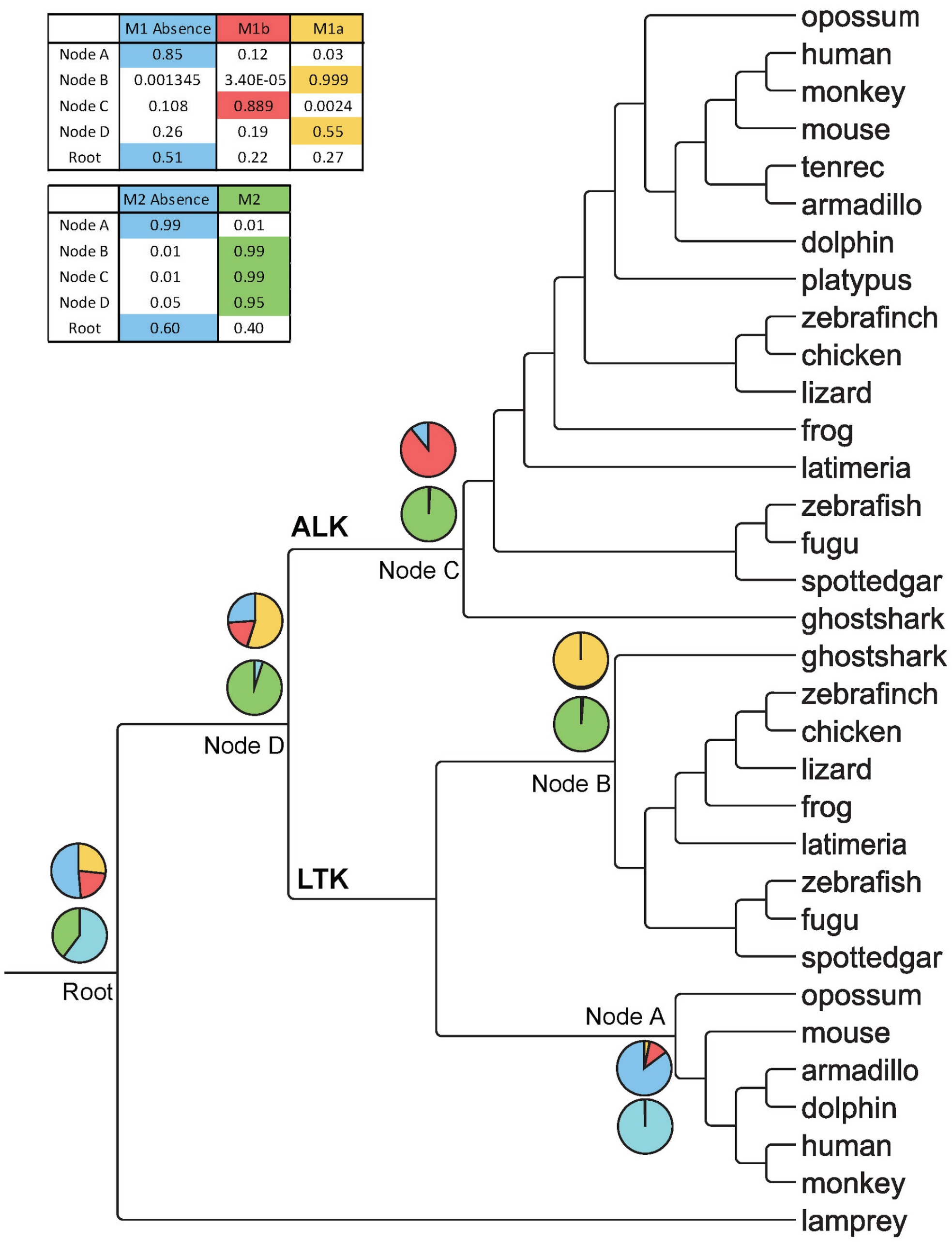


**Figure S2**. Estimated ancestral states of MAM domains for vertebrate ALK and LTK using maximum likelihood (ML) in BayesTraits (101, 102). MAM domains are are classified as MAM1a (M1a, red), MAM1b (M1b, yellow), and MAM2 (M2, green). Absence was coded in blue.

```
          Human        --GGKGGKNTMMRSHGVSVLGIFNLEKDDMLYILVGQQ-GEDACP-STNQLIQK---------VCIG  ENNVIEEE-IRVNR  HPERLENNSSVLGL
          Mouse        --GGKGGKNTMMRSHGVSVLGIFNLEKGDTLYILVGQQ-GEDACP-RANQLIQK---------VCVG  ENNVIEEE-IRVNR  HPERLESNSSVLGL
          Monkey       --GGKGGKNTMMRSHGVSVLGIFNLEKDDMLYILVGQQ-GEDACP-STNQLIQK---------VCIG  ENNVIEEE-IRVNR  HPERLENNSSVLGL
          Opossum      --GGKGGKNTMMRSHGVSVLGVFDLEKDDFLYILVGQQ-GEDACP-STSEIIQK---------VCIG  ENNVIEEE-IRANQ  HPERLENNSSVLGL
          Dolphin      --GGKGGKNTMMRSHGVSVLGIFNLEKDDVLYILVGQQ-GEDACP-STNQLIQK---------VCIG  ENNVIEEE-IRVNK  HPERLENNSSVLGL
          Platypus     --GGKGGKNTMMRSYGVSVLGIFDLQKGDTLYILVGQQ-GEDACP-HTNHVIQK---------VCIG  ESNVIEEE-IQVNR  HPERLENDSSVPGL
          Tenrec       --GGKGGKNTRMRSHGVSVLGIFNLEKDDMLYILVGQQ-GEDACP-STSQLIQK---------VCVG  ENNVIEEE-LRVNR  HAERLENNSSVLGL
          Armadillo    --GGKGGKNTMMRSHGVSVLGIFPLEKDDILYILVGQQ-GEDACP-SANQLIQK---------VCIG  ENNVIEEE-IRVNR  HPERLENNSSVLGL
          Zebrafinch   --GGKGGKNTMVRSHGVSVLGIFDLQKDDTLYILVGQQ-GEDACP-STNDVIQK---------VCIG  ENNVIEEE-IRVNR  HPERLENDSSIPGL
          Chicken      --GGKGGKNTMVRSHGVSVLGIFDLQKDDTLYILVGQQ-GEDACP-STNDVIQK---------VCIG  ENNVIEEE-IRVNR  HPERLENDSSVPGL
          Frog         --GGNGGKTPTL-AQGISLIGIFELQKDEILYILVGQK-GEDACP-RANSVLQM---------VCIG  ENTEIEDE-LRING  PPERLESNYTIPGM
          Lizard       --GGK-GKNTIMRSNGVNVAGIFKLQKNESLYILVGQQ-GEDACP-SANHIIQK---------VCLG  ENNVVEEE-IKANK  NFERLENDTSVPGL
          Latimeria    --GGRGVKSTMLRLHGMMVAGFFELQKDELLYILVGQQ-GEDACPHHTNPMIQK---------VCVG  ESNVIEEE-IRVNR  KPEKLETNTSIPGL
          Zebrafish    --GGRSV-LAVHKSHGVYMTGDFLLQKDELLYILVGQE-GEDACP-NMVPTMDR---------ICRE  QQGPS------INK  PEEVMDRDPSIPGR
          Spottedgar   --GGRSVEVT-ERSHGVYVVAEFLLQKDELLYILVGQQ-GEDACP-NPDPAFNQ---------LCLG  ESSLTD------NK  EEELLETDPSVPGQ
          Fugu         --GGRSV-LVMSRSHGVYLTGDFWLKKGEDLYILVGQQ-GEDACP-SSNNILNK---------ICQE  QSGPM------VNK  QLEQMDYELGQQGR
ALK       Ghostshark   --GGKGVKNDFKRFHGILVSATFELQKNEILYILVGQQ-GEDACP-NANPIIQK---------VCKG  ESNVIEEE-LRANG  FEEMLENDSFVPGL
------------------------------------------------------------------------------------------------------------------------
LTK       Human        --GGKGAKNHLSRAHGVFVSAIFSLGLGESLYILVGQQ-GEDACP-GGSPESQL---------VCLG  ESRAVEE-HAAMDG  SPEKLENRSEAPGS
          Monkey       --GGKGAKNHLSRAHGVFVSAIFSLGLGESLYILVGQQ-GEDACP-GGSPESQL---------VCLG  ESRAVEE-HAAMDG  SPEKLENRSEAPGS
          Mouse        --GGKGAQNHLSRAHGIFLSAVFFLRRGEPVYILVGQQ-GQDACP-GGSPESQL---------VCLG  ESG---E-HATTYG  VPERLETRAAAPGS
          Opossum      --GGKGAKNHNSRAHGIFVSAIFTLRQGELLYILVGQQ-GGDACP-GRTRESQD---------VCLG  ASSTTEEKEEKATG  SWEKLENQTSAPGT
          Dolphin      --GGKGAKDHLSPAHGVFLSAVFPLGHGELPYILVGQQ-EEDACP-KVSGATSS---------FCLG  ESPAAEE-HAATEG  TPEKLQNRSAAPGS
          Armadillo    RCGDKGAKSHVRRAXDLLVSSVFSLGCGEPLSTSGGSRAGLRSRS-RWSPRAAW---------RCLG  EPRAAEEP-AATEA  ARRQLEKRSAAPGA
          Zebrafinch   --GGKGAKNHNKRSHGVFISATFHLEKDELLYILVGQQ-GEDACP-GGNPETQK---------ICLG  ESSLIEED-YKKKR  PLEQFENSTAVPGV
          Chicken      --GGKGAKNHNERSHGVFISATFQLEKDELLYILVGQQ-GEDACP-GANPQTQK---------ICLG  ESSVIEED-YKKKK  PLEQFENSTAVPGV
          Frog         --GGKGAKNHNNPAHGVFISAIFPLQKDEILYILVGQQ-GEDACP-GKNELTKE---------ICMG  KSSIIEDN-YDDQN  HPERFENNTAVPGP
          Lizard       --GGKGAKNHNKRSHGVFISAIFKLEKDELLYILVGQQ-GEDACP-GASSQTMQ---------ICLG  ESSEIEEE-YKKEK  FLEQYVNNTAVPGI
          Latimeria    --GGKGAKNHNKRSHGVFISAIFHLEKDEILYILVGQQ-GEDACP-ARNAMTQK---------ICLG  ESSVIEDE-YTNNN  PLEQFENSTLVPGN
          Zebrafish    --GGKGAKNHNKRSHGVFISATFPLEKGDILYILIGHQ-GEDACP-GRNPQTHK---------ICLG  ESSVIEDG-FDSDG  FREQYENDTTVSGV
          Spottedgar   --GGKGAKNHNKRSHGVFISAVFPLEKGDILYILVGQQ-GEDACP-ARNPLTQQ---------VCLG  ESSVIEDDHFSGGS  PLEQYENDTTTSGA
          Fugu         --GGKGARNHNKRNHGIFISAIFPLEKGDVLYILVGHQ-GEDACP-GRNPETHK---------ICRG  GSSVIEDS---RQD  PLEQYTNSTKTPST
          Ghostshark   --GGKGAKNTNRRSHGVFISAIFHFEKGDNLYILVGQK-GEDACP-GKNPVTQK---------ICLG  ESSIIEDD-DHAND  LLEQYENDTLAPGV
------------------------------------------------------------------------------------------------------------------------
ALK       Lamprey      --GGKGARNNKQRAHGVYISANFILHKDEILYMLVGQQ-GGDACP-SVSSSCTSPQDHHCQRIVCRG  ESTVIEDE-KRKEG  VDDPFEGNLSVPGV
```

**Figure S3**. Analysis with DIVERGE (Gu et al. 2013) identified sites that exhibited significant substitution rate shifts among lineages between vertebrate ALKs and LTKs (green).

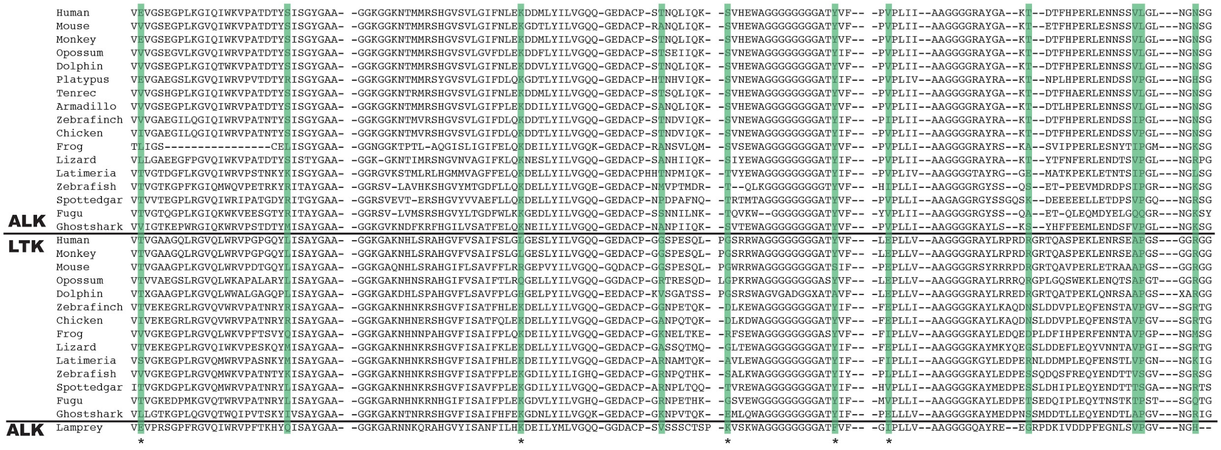


**Figure S4**. Analysis with `DIVERGE` (111) identified sites that exhibited significant substitution rate shifts among lineages within vertebrate LTKs (green).

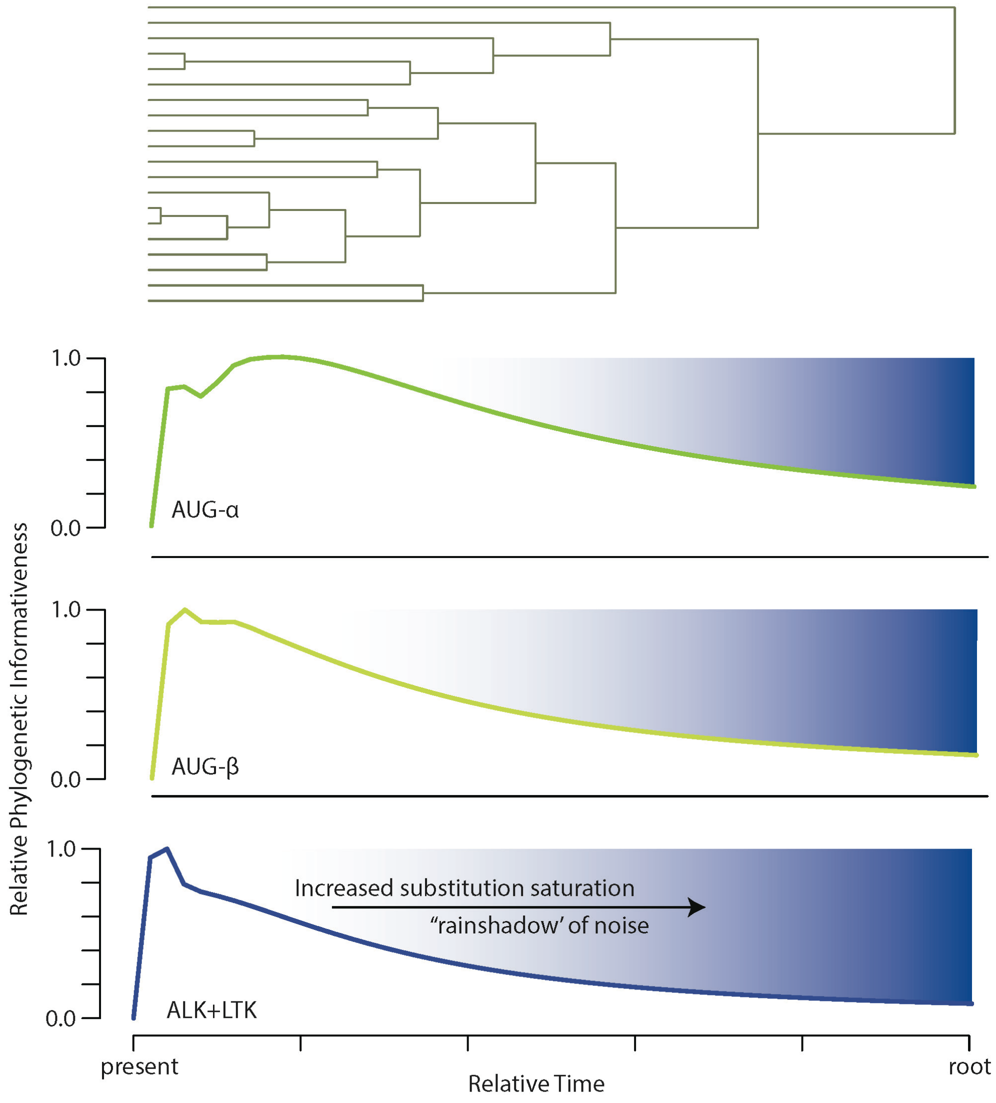


**Figure S5**. Phylogenetic informativeness profiles of AUG-α, AUG-β, and the clade comprising ALK+LTK lineages across vertebrate phylogeny (PhyInformR; 53). The decline in informativeness towards the origin of vertebrates indicates increased substitution saturation.

**Table S1**. Homologs of *ALK*, *LTK*, *JEB*, *HEN-1*, and *AUG* identified in sampled invertebrate and vertebrate genomes.

| Taxa | Common name | *ALK* | *LTK* | Ligand(s) |
|---|---|---|---|---|
| *Ailuropoda melanoleuca* | Panda | XP_019659786.1 | XP_019658000.1 | Aug-α ( ENSAMEG00000009559)<br>Aug-β (ENSAMEG00000015012) |
| *Anas platyrhynchos* | Duck | ENSAPLG00000013002 | ENSAPLG00000011988 | Aug-α (ENSAPLG00000012194) |
| *Anolis carolinensis* | Lizard | ENSACAP00000008904 | XP_008102570.1 | Aug-α (ENSACAG00000005420)<br>Aug-β (XP_008114420.2) |
| *Apis mellifera* | Honey bee | XP_016766222 | not found | Jeb (LOC100577600) |
| *Aplysia californica* | Sea hare | XP_012938087 | not found | Jeb-like (XP_005090980) |
| *Astyanax mexicanus* | Cave fish | ENSAMXG00000013686 | ENSAMXG00000011600 | Aug-α1 (ENSAMXG00000014141)<br>Aug-α2 (ENSAMXG00000008347) |
| *Bos taurus* | Cow | ENSBTAG00000007379 | XP_019823375 | Aug-α (XP_010799138.1)<br>Aug-β (XP_010810318.1) |
| *Branchiostoma floridae* | Lancelet | not found | not found | not found |
| *Caenorhabditis elegans* | Nematode | SCD-2 (WBGene00004740) | not found | Hen-1 (WBGene00001841) |
| *Callithrix jacchus* | Marmoset | ENSCJAG00000008763 | ENSCJAG00000020723 | Aug-α (ENSCJAG00000011653)<br>Aug-β (ENSCJAG00000014625) |
| *Callorhinchus milii* | Ghostshark | XP_007895095.1 | XP_007887733.1 | Aug-α (XP_007892194.1)<br>Aug-β (XP_007904100.1) |
| *Canis familiaris* | Dog | ENSCAFG00000005297 | XP_022268223.1 | Aug-β (XP_022268147.1) |
| *Cavia porcellus* | Guinea Pig | ENSCPOG00000008234 | XP_023423243.1 | Aug-α ( XP_003464974.1)<br>Aug-β (ENSCPOG00000032897) |
| *Chlorocebus sabaeus* | Vervet Monkey | ENSCSAG00000011035 | ENSCSAG00000007780 | Aug-α (ENSCSAG00000006838)<br>Aug-β (ENSCSAG00000014724) |
| *Choloepus hoffmanni* | Sloth | ENSCHOG00000012882 | not found | Aug-α (ENSCHOG00000011961)<br>Aug-β (ENSCHOG00000008251) |
| *Cimex lectularius* | Bed bug | XP_014247913 | not found | Jeb (LOC106666430) |
| *Ciona intestinalis* | Ciona | not found | not found | not found |
| *Ciona savignyi* | Ciona | not found | not found | not found |
| *Danio rerio* | Zebrafish | XP_691964.6 | ENSDARP00000061546 | Aug-α1 (ENSDARG00000091073)<br>Aug-α2 (ENSDARG00000042815)<br>Aug-β (ENSDARG00000074387) |
| *Daphnia pulex* | Water flea | EFX84662 | not found | Jeb (DAPPUDRAFT_263759, _264276) |
| *Dasypus novemcinctus* | Armadillo | XP_004452398.1 | XP_012376273.1 | Aug-α (ENSDNOG00000044907)<br>Aug-β (XP_004479910.1) |
| *Dipodomys ordii* | Kangaroo rat | XP_012869014.1 | XP_012865448.1 | Aug-β (XM_013009307.1.1) |
| *Drosophila melanogaster* | Fruit fly | FBgn0040505 | not found | Jeb (FBgn0086677) |
| *Echinops telfairi* | Tenrec | XP_012862293.1 | not found | Aug-α (NP_001002919.2) |
| *Eptatretus burgeri* | Hagfish | not found | not found | not found |
| *Equus caballus* | Horse | XP_023474203.1 | ENSECAG00000010591 | Aug-α (ENSECAG00000009187)<br>Aug-β (ENSECAG00000011169) |
| *Erinaceus europaeus* | Hedgehog | XP_007523966.1 | not found | Aug-α (XP_007529917.1)<br>Aug-β (XP_007516111.1) |
| *Felis catus* | Cat | XP_003984408.1 | confirmed* | Aug-α ( XP_023107938.1)<br>Aug-β (XP_023104497.1) |
| *Ficedula albicollis* | Flycatcher | ENSFALG00000011275 | ENSFALG00000006086 | Aug-α (ENSFALG00000002004) |
| *Gadus morhua* | Cod | ENSGMOG00000014962 | ENSGMOG00000011080 | Aug-α ENSGMOG00000012987 |
| *Gallus gallus* | Chicken | XP_015133623.1 | XP_004941681.1 | Aug-α (ENSGALG00000016364) |
| *Gasterosteus aculeatus* | Stickleback | ENSGACG00000012839 | ENSGACG00000008800 | Aug-α ENSGACG00000011449 |
| *Gorilla gorilla* | Gorilla | ENSGGOG00000004378 | ENSGGOG00000007006 | Aug-α (ENSGGOG00000014328)<br>Aug-β (ENSGGOG00000002382) |
| *Homo sapiens* | Human | NP_004295.2 | NP_002335.2 | Aug-α (NP_001002919.2)<br>Aug-β (NP_997296.1) |
| *Ictidomys tridecemlineatus* | Squirrel | ENSSTOG00000002811 | ENSSTOG00000008387 | Aug-α ( ENSSTOG00000021139)<br>Aug-β (ENSSTOG00000003032) |
| *Latimeria chalumnae* | Coelacanth | XM_014487828.1 | ENSLACP00000001197 | Aug-α (XP_014350854.1)<br>Aug-β (XP_006000473.1) |
| *Lepisosteus oculatus* | Spotted gar | XP_015208377.1 | XP_015194744.1 | Aug-α (XP_015213224.1)<br>Aug-β (XP_015208960.1) |
| *Loa loa* | Eye worm | EJD76771 | not found | Hen-1 (XP_003142890) |
| *Loxodonta africana* | Elephant | ENSLAFG00000014602 | ENSLAFG00000017768 | Aug-α (XP_023406155.1)<br>Aug-β (XP_010597850.1) |
| *Macaca mulatta* | Macaque | ENSMMUG00000010265 | ENSMMUG00000008170 | Aug-α (ENSMMUG00000017098)<br>Aug-β (ENSMMUG00000042459) |
| *Macropus eugenii* | Wallaby | ENSMEUG00000000874 | ENSMEUG00000002477 | Aug-α (ENSMEUG00000002734) |
| *Meleagris gallopavo* | Turkey | ENSMGAG00000005339 | XP_019471634.1 | Aug-α (ENSMGAG00000013966) |
| *Microcebus murinus* | Mouse Lemur | ENSMICG00000000637 | ENSMICG00000015875 | Aug-α (XP_012638705.1)<br>Aug-β (XP_012645510.1) |
| *Monodelphis domestica* | Opossum | XP_007476091.1 | XP_007480142.1 | Aug-α (ENSMODG00000014263)<br>Aug-β (ENSMODG00000025613) |
| *Mus musculus* | mouse | NP_031465.2 | NP_976220.2 | Aug-α (XP_017170402.1)<br>Aug-β (NP_001182661.1) |
| *Mustela putorius furo* | Ferret | ENSMPUG00000017981 | XP_012910813.1 | Aug-α ( XP_004745767.1)<br>Aug-β (XP_004777578.1) |
| *Myotis lucifugus* | Microbat | ENSMLUG00000010803 | XP_023613374.1 | Aug-α (ENSMLUG00000017708) |
| *Nomascus leucogenys* | Gibbon | XP_012356733.1 | XP_003266803.1 | Aug-α (ENSNLEG00000008389)<br>Aug-β (ENSNLEG00000000759) |
| *Ochotona princeps* | Pika | ENSOPRG00000007284 | ENSOPRG00000016526 | Aug-α ( XP_004582877.1)<br>Aug-β (XP_004580657.1) |
| *Oreochromis niloticus* | Tilapia | ENSONIG00000019537 | ENSONIG00000020096 | Aug-α1 (ENSONIG00000013462)<br>Aug-α2 (ENSONIG00000018754)<br>Aug-β (ENSONIG00000001740) |
| *Ornithorhynchus anatinus* | Platypus | XP_007665272.1 | not found | Aug-α (XP_003429395.2)<br>Aug-β (ENSOANG00000021848) |
| *Oryctolagus cuniculus* | Rabbit | ENSOCUG00000001085 | ENSOCUG00000026867 | Aug-α ( ENSOCUG00000029630)<br>Aug-β (ENSOCUG00000029695) |

| | | | | |
|---|---|---|---|---|
| *Oryzias latipes* | Medaka | XP_023807229.1 | ENSORLG00000015434 | Aug-α (ENSORLG00000013880)<br>Aug-β (ENSORLG00000005531) |
| *Otolemur garnettii* | Bushbaby | ENSOGAG00000014350 | ENSOGAG00000012850 | Aug-α (XP_012666040.1) |
| *Ovis aries* | Sheep | ENSOARG00000019617 | ENSOARG00000020370 | Aug-α (XP_014948423.1)<br>Aug-β (XP_014953145.1) |
| *Pan troglodytes* | Chimpanzee | ENSPTRG00000011796 | ENSPTRG00000006948 | Aug-α (ENSPTRG00000011606)<br>Aug-β (ENSPTRG00000024172) |
| *Papio anubis* | Olive baboon | ENSPANG00000015024 | ENSPANG00000018808 | Aug-α (ENSPANG00000015099)<br>Aug-β (ENSPANG00000015221) |
| *Pediculus humanus* | Body louse | XP_002432431 | not found | Jeb (XP_002425611) |
| *Pelodiscus sinensis* | Chinese softshell turtle | ENSPSIG00000010494 | ENSPSIG00000014814 | Aug-α (ENSPSIG00000002370)<br>Aug-β (ENSPSIG00000014916) |
| *Petromyzon marinus* | Atlantic Lamprey | ENSPMAG00000005264** | not found | Aug (annotated in this study)*** |
| *Poecilia formosa* | Amazon molly | ENSPFOG00000011525 | ENSPFOG00000005843 | Aug-β (ENSPFOG00000021789),<br>Aug-α1(ENSPFOG00000001067)<br>Aug-α2 (ENSPFOG00000022096) |
| *Pongo abelii* | Orangutan | ENSPPYG00000012522 | ENSPPYG00000006376 | Aug-α (ENSPPYG00000012693)<br>Aug-β (ENSPPYG00000018583) |
| *Procavia capensis* | Hyrax | ENSPCAG00000008629 | ENSPCAG00000005531 | Aug-α (ENSPCAG00000012119),<br>Aug-β (ENSPCAG00000007706) |
| *Pteropus vampyrus* | Megabat | ENSPVAG00000011977 | ENSPVAG00000016352 | Aug-α (ENSPVAG00000013389)<br>Aug-β (ENSPVAG00000005545) |
| *Rattus norvegicus* | Rat | ENSRNOG00000008683 | ENSRNOG00000025130 | Aug-α (ENSRNOG00000005154)<br>Aug-β (ENSRNOG00000048703) |
| *Sarcophilus harrisii* | Tasmanian devil | XP_023350365.1 | ENSSHAG00000017193 | Aug-α (ENSSHAG00000004800)<br>Aug-β (ENSSHAG00000009637) |
| *Sorex araneus* | Shrew | XP_004620066.1 | not found | Aug-α (XP_004616131.1),<br>(ENSSARG00000005846) |
| *Strongylocentrotus purpuratus* | Sea urchin | SPU_017036 | not found | Jeb-like (SPU_018625) |
| *Sus scrofa* | Pig | ENSSSCG00000008533 | ENSSSCG00000040414 | Aug-α (ENSSSCG00000039057)<br>Aug-β (ENSSSCG00000033922) |
| *Taeniopygia guttata* | Zebrafinch | XP_012427712.1 | XP_002199013.2 | Aug-α (ENSTGUG00000012951)<br>Aug-β (XP_004175323.1) |
| *Takifugu rubripes* | Fugu | ENSTRUP00000025266 | XP_011616029.1 | Aug-α (XP_011604626.1) |
| *Tarsius syrichta* | Tarsier | ENSTSYG00000009783 | ENSTSYG00000027024 | Aug-α (XP_008048005.1)<br>Aug-β (XP_008072903.1 ) |
| *Tetraodon nigroviridis* | Tetraodon | ENSTNIG00000008909 | ENSTNIG00000017323 | Aug-α (Q4RU86.1) |
| *Tribolium castaneum* | Red flour beetle | XP_008194070 | not found | Jeb (TcasGA2_TC001096) |
| *Tupaia belangeri* | Tree Shrew | ENSTBEG00000016366 | ENSTBEG00000006236 | not found |
| *Tursiops truncatus* | Dolphin | XP_019798032.1 | XP_019790652.1 | Aug-α (XP_004327977.2)<br>Aug-β (XP_004325501.1) |
| *Vicugna pacos* | Alpaca | ENSVPAG00000004035 | XP_015092567.1 | Aug-α (XP_015106603.1)<br>Aug-β (ENSVPAG00000010782) |
| *Xenopus tropicalis* | Frog | XP_017949451.1 | XP_002933782.2 | Aug-α (XP_012818929.1)<br>Aug-β (XP_012820744.1) |
| *Xiphophorus maculatus* | Platyfish | ENSXMAG00000009140 | ENSXMAG00000013305 | Aug-α (ENSXMAG00000007504)<br>Aug-β (ENSXMAG00000018426) |

* Annotation of the presence of an ALK homolog in this genome is confirmed by NCBI in their current release; however, sequences are not publicly available.

### ** Annotation of ALK in lamprey genomes:

>>*Lethenteron camtschaticum* (Japanese lamprey ALK predicted with intron positions (size in bases)

TPLTPHLRLSVKDNMLQWMFKTCGASGPLGPTQVQCDNAYRNTNITVEVPRSGPFRGVQIWRVPFTKHYQ(1767)I
SAYGAAGGKGARNNKQRAHGVYISANFILHKDEILYMLVGQQGGDACPSVSSSCTSPQDHHCQRIV(5354)CRGES
TVIEDEKRKEGKVSKWAGGGGGGGGATFVFKT(7787)GYGIPLLVAAGGGGQAYREEGRPDKIVDDPFEGNLSVPG
VNGHTG(3023)GGGGWYGHVGPARSGRSLNQGGQGGSACPEALVNWAWATSGGFGGGGGACSAGGGGGGYIGER(1
263)DSAFDNNIMADGESGVSFMHPDGGLFTMPLAI(4228)MERDGEVVVEEYLNCSHCESKLCKRNADEHDSSIV
CHCQEGVLASDGFSCSVTTRPPRD(4166)PKNVFHLLVSVLEPFSFCIIKC(3775)CVVYRRKQAQLQAVRMELL
SPEYKLSKIRTSTIMTDYNPNYCFAGSPAALTDLKEVPRRNINLLR(2737)ALGHGAFGEVYEGRVTGIEPSPLKV
AVKT(1375)LPEICSEQDELDFLMEALIIS(4146)KFNHHNIVRCVGVSLQTLPRFILLELMAGGDMKTYLREER
PRAVCAGPQK(867)MLDLLTMAKDIACGSQYLEDNHFIHR(2118)DIAARNCLLTSKGSDRIAKIGDFGMARDIY
R(3635)ASYYRKGGRAMLPVKWMPPEAFLEGIFTSKTDTW(2764)SFGVLLWEIFSLGYMPYPSKSNQEVMEFVT
SGGRMDPPKNCPVPV(570)YRIMTQCWQHRPEDRPNFSTILERITYCTQD(741)PDVVQTPLPTEYMPPPDDEGN
TVMRPPNPDRVPPLPMSGPLEAAPFAAPPQPARP(188)ANGNNLWNPTYGS

>*Petromyzon marinus* (Atlantic lamprey) ALK predicted (Partial)

PPSPLTPHPRLSAKDNTLQWMFKTCGASGPLGPTQVQCDNAYRNTNITVEVPRTGPFRGVQIWHVPFTKHYQISAYG
AAGGKGARNNKQRAHGVYISANFILHKDEILYMLVGQQGGDACPSGSSSRIGQGVCRGESTVIEDEKRKEGKVSKWA
GGGGGGGGATFVFKMQDGVRIPLLVAAGGGGQAYREEGRPDKIVDDPFEGNLSVPGVNGHTGAAGGGGGWYGHVGPA
RSGRSLNQGGQGGSACPEALVNWAWATSGGFGGGGGACSAGGGGGGYIGGNSASDNNIMADGESGVSFMHPDGGLFT
MPLAIMERDGEVVVEEYLNCSHCESKLCKRNADEHDSIVCHCQEGVLASDGFSCSVTTRPPRVFHLLVSVLEPFSFF
IIKVQRCCVVYRRKQAQLQAVRMELLSPEYKLSKIRTSTIMTDYNPNYCFAGSPAALSDLKEVPRRNINLLRALGHG

AFGEVYEGRVTGIEPSPLKVAVKTLPEICSEQDELDFLMEALIISKFNHHNIVRCVGVSLQTLPRFILLELMAGGDM
KTYLREERPRAVCAGPRKSMLDLLTMAKDIACGQYLEDNHFIHRDIAARNCLLTSKGSDRIAKIGDFGMARDIYSRA
SYYRKGGRAMLPVKWMPPEAFLEGIFTSKTDTWSFGVLLWEIFSLGYMPYPSKSNQEVMEFVTSGGRMDPPKNCPVP
VYRIMTQCWQHRPEDRPNFSTILERITYCTQDPDVVQTPLPTEYMPPPDDEGNTVMRPPNPDRVPPLPMSGPLEAAP
FAAPPQPARPANGNNLWNPTYGS

>Contigs identified with reads that were mapped to regions of the genome between predicted ALK exons.

**ALK_intron_contig_1**
ACGTCACGCACCTGGGGGGGGGGGTCAGACATTTGTGACGATGTGTGACGAGGGGGGGGGT
TGAGAAATGTGACGTCACATAAAAATGCCCAACTTTAAATACATATAGACAACACGTTCTAC
TTAATAAAATAGACTTTTTAATAAATGTTTTCTCCTACAACATCAGAGTGCAGAGACACATT
AAATACGCACTATAATGACAATAGTTTAAAGCAATTTGAAGCATGTTTTATTTTCAAAATGG
AAATATGGTTTTGGGGGGGGGGGCAGAGCTTTGTGACGTCATATTTGAGGGAGGGTCTGACT
TTTTGTGACG

BLAST search against NCBI nucleotide database (the best 2 scored):
>>Lethenteron camtschaticum clone BAC 256N19, BAC 343N18, BAC 232I2 homeobox protein Hox-alpha9 (Hox-alpha9), homeobox protein Hox-alpha8 (Hox-alpha8), homeobox protein Hox-alpha7 (Hox-alpha7), homeobox protein Hox-alpha6 (Hox-alpha6), and homeobox protein Hox-alpha5 (Hox-alpha5) genes, complete cds; miR-10 miRNA gene, complete sequence; and homeobox protein Hox-alpha4 (Hox-alpha4) gene, complete cds Sequence ID: KF318001.1 Length: 138871 Number of Matches: 2 (89%).

>>Lethenteron camtschaticum clone BAC 238G18 homeobox protein Hox3-alpha (Hox-alpha3), homeobox protein Hox2-alpha (Hox-alpha2), and metaxin alpha (Mtx) genes, complete cds, complete sequence Sequence ID: KF318005.1 Length: 94233 Number of Matches: 2 (88%).

**ALK_intron_contig_2**
AAGCTCTCTCTGTGTGAAAAAAGCTAGCCCGCATCAATACCTTTTAAGCAAACTGGGCCTGC
TGAAATACAAACCTGTCAACCCCTTATCAGTGCCTACTTTATTACAGACCAATTCAGACCTTA
TTGGTCACCCCGTGCCCTTATAAATCTCAACGTTCATCCTACAAAAGTCTCATCACAAGAGA
GAAACACAAACAAATA

BLAST search against NCBI nucleotide database (the best 2 scored):
>>Lethenteron camtschaticum MYH2, MYH3, MYH1 genes for myosin heavy chain 2, myosin heavy chain 3, myosin heavy chain 1, complete cds, Sequence ID: AB720829.1 Length: 106535 Number of Matches: 2 (97%).

>>Lethenteron sp. N DNA, microsatellite loci Lspn 013 Sequence ID: AB209974.1 Length: 579 Number of Matches: 1 (89%).

**ALK_intron_contig_3**
TGACTGCAGGAATTACTCTTTGGTTTTTTCTGTAAAGTCACGTATAAATATGAAATAATATAA
TACCTTAATAGCGTACGACGTTTCGGTTGCAACACAAGCAGCCATCATCAGGTACAGAAGAG
CTAAATTTACGAGAGAGAGGAAACACATAACCCCAGATGTATATACAGACACCCAAGTAGG
GGGAACTAACACAGATGCAACAGGGGACACAGAGAGAGAAGCATACCAAACAAACAGGGC
TTAGCAAAATTTGATCAAATTACTCGT

BLAST search against NCBI nucleotide database (the best 2 scored):
>> Petromyzon marinus lamprin L-1.8 (L-1.8) gene, complete cds, alternatively spliced
Sequence ID: AH007080.2 Length: 8277 Number of Matches: 1 (92%).

>> Petromyzon marinus lamprin L-0.9 (L-0.9) gene, complete cds, alternatively spliced
Sequence ID: AH007079.2 Length: 6543 Number of Matches: 1 (89%).

**ALK_intron_contig_4**
GCAGAAAGCGACCCGTCCCAGCACACCGGCATGCCGCTGTCGCTGGTGCTGTCCGTGGTGAC
ACTGGCAACGGCCGCTGTGCTTGTGCTCACCTGCTCGGCCATCATGATCG

BLAST search against NCBI nucleotide database:
>> No significant similarity found.

**ALK_intron_contig_5**
TGTCTCCCCAGTGGCGATGTTTACACACGACACCAGGTACTCATTTGAGGCTGAGTCGACCT
AGGGGAGGCCCAGGACCGGTTCAAATAATCGTCACTGGTGTTGTCCATGAGCGGGAATTGA
ACTTAGGTCGCCGGGTTCGTAGCGTTGCGC

BLAST search against NCBI nucleotide database (the best 2 scored):
>> Lethenteron camtschaticum clone BAC 193M22 homeobox protein Hox-epsilon4 (Hox-epsilon4) gene, complete cds; and homeobox protein Hox-epsilon1-like (Hox-epsilon1) gene, partial sequence, complete sequence; Sequence ID: KF318007.1 Length: 44119 Number of Matches: 10 (92%).

>> Lethenteron camtschaticum MYH2, MYH3, MYH1 genes for myosin heavy chain 2, myosin heavy chain 3, myosin heavy chain 1, complete cds; Sequence ID: AB720829.1 Length: 106535 Number of Matches: 1 (91%).

#### *** Annotation of AUG in lamprey eel genomes:

>*Lethenteron camtschaticum* (Japanese lamprey AUG predicted with intron positions (size in bases)
MAAIRVHALLGLLALLATSARCWDGAAGLSSRQSLLDLVLAALEELERGGGGTRHSSSPLAFPSSEDLAGNAAEMSR
IPSEIFFPI(2671)AIDTRDPSLKEKFIKHVTNK(2977)SSMYMSPECSRQFYRLYHNTRDCTVRSCKSALLHGS
DINP-

>*Petromyzon marinus* (Atlantic lamprey) AUG predicted
MAAIRVHALLGLLALLAASARCWDGAAGLQAGLASRQSLLDLVLAALEELERGGGTRHSSSPLAFPSSEDLAGNAAE
MSRIPSEIFFPIAIDTRDPSLKEKFIKHVTNKSSMYMSPECSRQFYRLYHNTRDCTVRSCKSALLHGSDI-

>Contigs identified with reads being mapped to the genome regions between predicted AUG exons.

**AUG_intron_contig_1**
GAGGGAGAAAGAAAGCAAAACACGAGGCCACCGTTAGCTCACGGCCAGTGCCGATCGCTCCCGA
CCGCGAGCTGCCCGCACTGCCACTGGCAGACAGACGCACACATCGAGAGGGACAACGAACACTG
CGGGAATTTTATTTCACGCGCACGCATAACCCTGCTAATCGCACCTCCGATTAGCACGGTTGGCT
ACGTACGGCGCGACAAAAGTCCAACGTACGTAGATAAGGAGATTTTAAACATTCACACATTACA
ATCAGCGAGCAGTTGGTGGAGGAAGCTCCGCAGAGCCCCTGCTGGAGGCTATTTTGGCCTACTTT
TTCAAACTCGCCAGATGTCTCGAGAGACGTGAGCTTGCCCACACTTTACCCCGCCGACACACGAC
GTGCCCTGGGTAGAGGCGGCGGGGTTTGGTAGCGGGGCGTTCGAGAAACTCGCACCACCGTTTA
AAAGCCGCCGCGCCGGGCGATGGAATCGAACCGGTGACTTCTTGGGTCCTGCGCGTCGTGCACGC
TGACCACCGCACCGCGGGCGCGGCTCCGGAGGGAAGGGGCCGTGGAGCCCAACGTGAGGCGCTC
TTTGAGCGTCGTTGATCGGCGCTGGGCCCGTAAGGATTGGTCAGTCGCGAAAGTCGCGAGCGCCC
CCAGATTTCATTGGATTTTGACGTTCGCGCGCTGCATATGTACGGGGCGATCCATGTATAGACAC
CCCATTGATTATAATCGTCAATGAACTACATTCTGAACAGGTGGTTGCAGACTCTGAAAGCGGTG

GTTCCTTGGCTGACGGAAGAAACACCTTTACTATTGAATAATACGTGCGCTGAGATTGGTGGATG
TTTCTACATATATGTTTTGAAATAACAATGTATTTTCTTCACAACGTATCCCTGTTGTTTCAATGCG
GTATCTATAAATAGATCACCGTCCAGCTTCTGAAGCAACCATTTGATTGACACGAAACGTCGTGA
GAAGGGATGGACGATGGCCGATGGCGAACAATAACAACGGAGTTGCGGCCAAGGAATGAAACG
TCCACGGCTGAGGTGGAGCGATGAAGTTCGTGGACGATCATGCCCGAAACGATCGTAAAGTGGT
AGGCTTGGGGGGAGGCCTTCATCCAGCAGCGGATTGACCGTGGCCGATGATGACGGAGATAATT
TGCGGCCACGCTCCTGGGTAAGCAGTGACGAGGCACGGGGAGCAACCGGGGAACTTTCGTCACC
CCCCCCCTCCGTTCGGTGGTGGGCGGCTTTGTACCGGGCTGCTCGCGTTGAGTCGTCATACTAACA
GTTCTTTTTACATAAAAAAAGACACGGGGAGCTGCCGCATCCACCGCACGGTGGGCCCTCGGCG
CTCGTTGGGGCCTCACGCGGAGCAACGCGGCGGGGTCGCGAGTGAAGGGTCGCTGGGTCACACG
GGGCCACGTGTTCAACAGGGCTTCCCGCGTGGCGCCTCTCCTCCCTGTGTCACCGCGAGCGATGG
ACCGTGGCCAGCCAAGATTCAAAATTACAGAAATGACTCCCCCCGCCCCCGCTGGGATGACTTTT
TGGCCGCCGCGAGGATGAACGTTGGTGACATCACGCCCGAGGTGGAGTGTTGCTTGGGCAAGGT
CCAGGACGCACGCGTTATGATGGCGGGTATCGATAGAACGCATTGGCAGACAATGTCAGGCAAG
TGGCGTTATTAAATGCCAACTAAGGCTGCCTGCGAGGAGGTAGTGTATGGCTATCATGTCCGCCG
CATGAGATGCTATTTATTTTGGCCAAATCAACCTAGGAGTGGGCCAGGTAACAAAAACATCCGG
CAGGTGCGCAAGCCCCAGAATTGAATTAGTTTTCAGTTTCATTTATTTGGTTTAACCCTGTGAGC
CATTCTGCTCTTGATTCGGGTGCAGCTGCAACAAACAATAAAATTACGAGATACATTTCAAATTT
ACTGCACGCAAATTGAAACATCCCCTATAAAAAAAATTACATCTTCCAAAAAAAGCACCGTATGC
TCATACTGCGTAAAAAAATGTGCAAAAATCCCAAGAATCCGGAACGATGATTCCGTTCGCAGGA
GATCAAAATCAAAATACACATGCACACATATACATAAACACAAACACTGCGATTCTCCGTCTAGC
GTTAATGACCGTTGTAACAAGTGCCGTTAGCTGTATCGACCTTCATGAGGTCGATTCAGCGGCAT
GAAGGAGAGGTGTGACCAGATCCAAGCCCTACCGCCGTTGTCTCTCCGGTGCTGACATTTACACA
CCACACCAGGTACACGTGTTTCATGGTGAGTCGACCAAGGGCAGACTCGCGACCGGTTCAGATAT
TCGTCACCAATGTTGACCGTTGCTGGGAAATCGACCTCTGGTCGTCTGGTTCGTAGCGCGAGGCA
TTTTCCATTACATTCCTACTCCATAATCACGCACGCACTCGCAAGAGACAAAGATCCTCCCCGAT
AGACTTAACAGACTTTTTGGGGGGATAGCGATGTAAGCGAGCACCTATGAAGGCCGGCACGGCA
CACACGAAGGGGTGGGTGGCCAACACACACGCTGACACACACACACACACTGACACACACACAC
ACACTGACACACACACACACTGACACACACACACACTGACACACACACACTGACA

BLAST search against NCBI nucleotide database (the best 2 scored):
>>Lethenteron camtschaticum clone BAC 424C19, BAC 204J11 homeobox protein Hox-epsilon14 (Hox-epsilon14) gene, partial cds; and homeobox protein Hox-epsilon13 (Hox-epsilon13) and homeobox protein Hox-epsilon11 (Hox-epsilon11) genes, complete cds, complete sequence; Sequence ID: KF318004.1 (83%).
>>Lethenteron camtschaticum clone BAC 238J5 homeobox protein Hox-delta3 (Hox-delta3) and homeobox protein Hox-delta2 (Hox-delta2) genes, complete cds, complete sequence; Sequence ID: KF318008.1 (84%).

**AUG_intron_contig_2**
CGAATCCTATAGAACGGCCGATATTTAATCCACGCAGAAACGATCAAGGATTAATAATATTAACG
TGTTTAAGCAAACAGATAGTCCACCGCCGGACTATCTAACTCATTTGAAGGCCAATGTAGACAAA
TTATTATCGAGGCTTCA

BLAST search against NCBI nucleotide database (the best 2 scored):
>>Lethenteron camtschaticum MYH2, MYH3, MYH1 genes for myosin heavy chain 2, myosin heavy chain 3, myosin heavy chain 1, complete cds; Sequence ID: AB720829.1 (84%).
>>Lethenteron camtschaticum clone BAC 256N19, BAC 343N18, BAC 232I2 homeobox protein Hox-alpha9 (Hox-alpha9), homeobox protein Hox-alpha8 (Hox-alpha8), homeobox protein Hox-alpha7 (Hox-alpha7), homeobox protein Hox-alpha6 (Hox-alpha6), and homeobox protein Hox-alpha5 (Hox-alpha5) genes, complete cds; miR-10 miRNA gene, complete sequence; and homeobox protein Hox-alpha4 (Hox-alpha4) gene, complete cds; Sequence ID: KF318001.1 (90%).

**AUG_intron_contig_3**
GCTGTCTGGGTGGTGGTGGTGGCGGTTCAAGGTAGCGTGGGAGAATGTGGGCGTGTGTGGGCAA
GTGGCGTGGCGGGGAAGGTGGGGGTCGAACGGTGTTACATTGTATAATGGGGTATTCGTTACCTT
TGTGCGCTATATTATTATTATTATTGCTGGTTCTCTAGTGTCCCTGAAGCACCCCGCAACCCTCGT
TTGGATAAGAGGCAATTTCAGGGACCTCATTTGTAGTTCTGTGAGATAATCTTTCTAGAGGCCGC
TCTTCAGGAGATTCATTACACTAAACTATGTTTTATTTTACCGGGTAAGGCCCATTTAGCTATATG
GCTGTTTTTTTCAGAAGCGACCTGGCCACAAATGATAGCCCAATGATGTGGCTTGTCACTTTGAA
ATCTATTGCAGCCAAATACTCTAAACGCATGTTCATTCTAAATGTGGAGGGAAAAACCAGGAGA
AAGATCCACGTGAACATGGGGAGAGAGAACATGCAAACTCTAAATATTCAGACATCAGGATTAG
GATCAAACCCAAGACCCCCTGCATACCAGGCAAGGTAGTTAAGATCTCGCCACCGCGGCCTCCCA
TCAGCTACGTAGGTTTTCCTGGGGTCGGTGAACCAGACATTATCTCAGTGACCTACACAGTATGT
CTCTCACAATGGTTCTGCCTACTACTAGTCAAATGCCACCTTACTTCAGTCTACACCAGGGATCTG
GAAAATTATTCCGTTTGAAATGTT

BLAST search against NCBI nucleotide database (the best 2 scored):
>>Petromyzon marinus ABCB9-like TAP-family protein (ABCB9) gene, complete cds; Sequence ID: AH012171.2 (79%)
>>Lethenteron camtschaticum clone BAC 424C19, BAC 204J11 homeobox protein Hox-epsilon14 (Hox-epsilon14) gene, partial cds; and homeobox protein Hox-epsilon13 (Hox-epsilon13) and homeobox protein Hox-epsilon11 (Hox-epsilon11) genes, complete cds, complete sequence; Sequence ID: KF318004.1 (78%)

**AUG_intron_contig_4**
CAAAAAATGAGTGTTGGTGGTAAGTGAATTAGCAAGAGAGAGTGAGCGGGCGAGACAGTCAAG
AGGGAATGAGTGAGCGAGGAAGCGAGTGACCAACAGGGAATGAGCGAGCCAGTCAGTGAGTGA
GTAAGTGAGGGGGCGAGTGACCAAGAGGGAATGAGCGAGCCAGTCAGTGAGTGAGCAAGAGTG
TGTGGGCAAGTGAGAGAGCGAGTGAGCAAGAGGGAGTGAGCGAGCGAGTCAGTGAATTAGTGA
GTGAGGGAGCGATTGGGCTAGAGGGAGAGGGAGTGAGCGAACGAGGGAGTGAGCAATCACGCG
GGGGTGAAAACTGTGA

BLAST search against NCBI nucleotide database:
>> No significant similarity found

**AUG_intron_contig_5**
CGATGTCGAGCCCCCACCCCGCCCCCCTCCCACCGGTGTTCGCGCACTCCGAGGGATTCATCTCA
TCCAGGGCTGTGCGGCACAGCGAGCGCGGAACGAAAATACCTCCAAACCCTGCACACCCACATCC
AGTATCTCGCGCTATGGCCGTGGCGACTACAAGTCACATGCCACTTGCTGTAGTCTGGCAGCAAC
CCGGTATGGATTACATGACCAATTATGCCTGGCGCATTCCAACAACAACTATAATAATGATGTCC
TTTATTTAGCCACGAAAGCGTCACTGAGTTCCCAGACTCTTTTCTATCAGGAGGTTTCCTGCAGTG
GGATGTTCGTGTCCATAGGGGAAGGAGGCGACAGTGCTTGGAGAAAACCCATCACACATACCGC
GGGGGAAACACTCAAAATTCCACAAGAATAGATTCAAGAATCTCCTGGCCACCATGTCACCAAG
TGTGCGGTGGTGACACGTGAGGCTTTGAGTCTGGATCTGTGCGCGTGGTGTGTATACTGTAACAA
AGACATCAATATTACAGTCACAAAGGCTGATGCGGCAAGAGCTGTTAGCGAGCACCTATTAAGG
CCGGCACAGAGCACACCAAGGGGGTGGTGTGGGCAAGTACCACACCCGCCCGCCTTTGTCTCCCC
ATTTCTCATGTTTACACACAAAACCTTGTCCTCGTTTAACTCTGAGTCGACTTAGGGACCGGTTAA
GATATTTTACCGATGATGTTGGCCAAGTCCGGGAATCAAATCAACCTGGGGTCACTGGGTTCATG
GCGCATAGCGCAGTGCGCTAACCACTGCGCTACT

BLAST search against NCBI nucleotide database (the best 2 scored):
>>Lampetra fluviatilis partial mRNA for neurotrophin type 2 (nt2 gene); Sequence ID: FR693754.1 (80%)

>>Lethenteron camtschaticum clone BAC 424C19, BAC 204J11 homeobox protein Hox-epsilon14 (Hox-epsilon14) gene, partial cds; and homeobox protein Hox-epsilon13 (Hox-epsilon13) and homeobox protein Hox-epsilon11 (Hox-epsilon11) genes, complete cds, complete sequence; Sequence ID: KF318004.1 (87%)

**AUG_intron_contig_6**
AATCCACGCAGAAACGATCAAGGATTAATAATATCCACCGCCCGACTATCTAACTCATTTGAAGG
CCAATGTAGACAAATTATTATCGAGGCTTCAC

BLAST search against NCBI nucleotide database(the best scored):
>> Lethenteron camtschaticum MYH2, MYH3, MYH1 genes for myosin heavy chain 2, myosin heavy chain 3, myosin heavy chain 1, complete cds; Sequence ID: AB720829.1 (97%)

**AUG_intron_contig_7**
ACACGCAATAAGGGGCCATCCATTTAGTAGGTACGCAAAAAATCTGTCAAAACTTGACCCCTCCC
CC

BLAST search against NCBI nucleotide database:
>> No significant similarity found

**AUG_intron_contig_8**
AAAGTGGAAATGTTTATATTGACAATATGATGCCAAGAGCACAGTACAAATGTAACAGATAAGC
AAATGCATAAAAGCAAAAACCAAAGCCTCGTTTT

BLAST search against NCBI nucleotide database:
>> No significant similarity found

**Table S2**. Gene-specific primers designed for RACE on lamprey AUG.

| Identifier | Sequence[a] |
|---|---|
| **GSP #1** | GATTACGCCAAGCTTccatacgcgtgcacgcgctgctgg |
| **GSP #2** | GATTACGCCAAGCTTcgagatattcttccccatagcg |
| **GSP #3** | GATTACGCCAAGCTTccaataagagttccatgtacatgtcaccg |
| **GSP #4** | GATTACGCCAAGCTTgctgctgggacggagctgcgggtctc |
| **GSP #5** | GATTACGCCAAGCTTcgcattcccctcctccgaggacctcgc |
| **GSP #6** | GATTACGCCAAGCTTgcagttctaccggctctaccacaacacccgcg |

[a] 15 bases of anchor primer (upper case) and gene specific primer (lower case).

**Table S3**. Sea lamprey transcriptomic data from NCBI analyzed to evaluate expression of *ALK* and *AUG*.

| Run # | *ALK* | *AUG* | BioProject | Size (MB) | Description |
|---|---|---|---|---|---|
| SRR068400 | + | - | PRJNA132763 | 56 | olfactory epithelia |
| SRR068401 | - | - | PRJNA132763 | 46 | olfactory epithelia |
| SRR2238808 | + | + | PRJNA294488 | 21783 | embryo stages 23–25 |
| SRR2238818 | + | + | PRJNA294488 | 21590 | embryo stages 26–28 |
| SRR3002835 | + | + | PRJNA306044 | 537 | 1 day post-fertilization embryos |
| SRR3002836 | + | - | PRJNA306044 | 543 | 1 day post-fertilization embryos |
| SRR3002837 | + | + | PRJNA306044 | 3050 | 1 day post-fertilization embryos |
| SRR3002838 | + | - | PRJNA306044 | 471 | 2 day post-fertilization embryos |
| SRR3002839 | + | + | PRJNA306044 | 475 | 2 day post-fertilization embryos |
| SRR3002840 | + | + | PRJNA306044 | 2445 | 2 day post-fertilization embryos |
| SRR3002841 | + | + | PRJNA306044 | 502 | 2.5 day post-fertilization embryos |
| SRR3002842 | + | - | PRJNA306044 | 506 | 2.5 day post-fertilization embryos |
| SRR3002843 | + | + | PRJNA306044 | 2537 | 2.5 day post-fertilization embryos |
| SRR3002844 | + | - | PRJNA306044 | 568 | 3 days post-fertilization embryos |
| SRR3002845 | + | - | PRJNA306044 | 572 | 3 days post-fertilization embryos |
| SRR3002846 | + | - | PRJNA306044 | 2943 | 3 days post-fertilization embryos |
| SRR3002847 | + | - | PRJNA306044 | 573 | 4 days post-fertilization embryos |
| SRR3002848 | + | - | PRJNA306044 | 577 | 4 days post-fertilization embryos |
| SRR3002849 | + | - | PRJNA306044 | 2819 | 4 days post-fertilization embryos |
| SRR3002850 | + | - | PRJNA306044 | 424 | 5 days post-fertilization embryos |
| SRR3002851 | + | - | PRJNA306044 | 428 | 5 days post-fertilization embryos |
| SRR3002852 | + | + | PRJNA306044 | 2198 | 5 days post-fertilization embryos |
| SRR369904 | + | + | N/A | 2533 | meiotic testes |
| SRR388671 | + | - | PRJNA50489 | 15 | Parasitic Olfactory Epithelium |
| SRR388684 | + | - | PRJNA50489 | 19 | Adult Olfactory Epithelium |
| SRR388685 | + | - | PRJNA50489 | 17 | Adult Brain |
| SRR388686 | + | - | PRJNA50489 | 16 | Larval/Parasitic Brain |
| SRR388687 | - | - | PRJNA50489 | 14 | Larval Liver |
| SRR388688 | - | - | PRJNA50489 | 13 | Parasitic liver |
| SRR388689 | + | - | PRJNA50489 | 1302 | Larval intestine |
| SRR388690 | + | - | PRJNA50489 | 1311 | Larval Kidney |
| SRR388691 | + | + | PRJNA50489 | 1368 | Small Parasitic Kidney |
| SRR388692 | + | - | PRJNA50489 | 1527 | Embryo (Late Blastula, Stage 18) |
| SRR388693 | + | - | PRJNA50489 | 1765 | Embryo (Gastrula, Stage 20) |
| SRR388694 | + | - | PRJNA50489 | 1849 | Embryo (Neurula, Stage 22a) |
| SRR389301 | + | + | PRJNA50489 | 1938 | Adult Brain |
| SRR389302 | + | + | PRJNA50489 | 1801 | Parasitic Liver |
| SRR389303 | + | - | PRJNA50489 | 1288 | Small Parasitic Proximal Intestine |

| | | | | | |
|---|---|---|---|---|---|
| SRR389304 | + | - | PRJNA50489 | 1206 | Small Parasitic Distal Intestine |
| SRR389306 | + | + | PRJNA50489 | 1240 | Adult Intestine |
| SRR389307 | + | - | PRJNA50489 | 1249 | Adult Kidney |
| SRR389308 | + | + | PRJNA50489 | 1909 | Embryo (Neurula, Stage 22b) |
| SRR389309 | + | - | PRJNA50489 | 1843 | Embryo (Neural Crest Migration, Stage 23) |
| SRR389310 | + | - | PRJNA50489 | 1832 | Embryo (Neural Crest Migration, Stage 24c1) |
| SRR389311 | + | + | PRJNA50489 | 1763 | Embryo (Neural Crest Migration, Stage 24c2) |
| SRR5124568 | + | + | PRJNA358429 | 3213 | olfactory tissue after 24 hours exposure to 5g/mL of copper |
| SRR5124569 | + | + | PRJNA358429 | 2940 | olfactory tissue after 24 hours exposure to 5g/mL of copper |
| SRR5124570 | + | + | PRJNA358429 | 2136 | olfactory tissue after 24 hours exposure to 30 g/mL of copper |
| SRR5124571 | + | + | PRJNA358429 | 3158 | olfactory tissue after 24 hours exposure to 10 g/mL of copper |
| SRR5124572 | + | + | PRJNA358429 | 2709 | olfactory tissue after 24 hours exposure to 30 g/mL of copper |
| SRR5124573 | + | + | PRJNA358429 | 2687 | olfactory tissue after 24 hours exposure to 0 g/mL of copper |
| SRR5124574 | + | + | PRJNA358429 | 2442 | olfactory tissue after 24 hours exposure to 5 g/mL of copper |
| SRR5124575 | + | + | PRJNA358429 | 2770 | olfactory tissue after 24 hours exposure to 5 g/mL of copper |
| SRR5124576 | + | + | PRJNA358429 | 3169 | olfactory tissue after 24 hours exposure to 0 g/mL of copper |
| SRR5124577 | + | + | PRJNA358429 | 2651 | olfactory tissue after 24 hours exposure to 0 g/mL of copper |
| SRR5124578 | + | + | PRJNA358429 | 2621 | olfactory tissue after 24 hours exposure to 30 g/mL of copper |
| SRR5124579 | + | + | PRJNA358429 | 2511 | olfactory tissue after 24 hours exposure to 30 g/mL of copper |
| SRR5124580 | + | + | PRJNA358429 | 2748 | olfactory tissue after 24 hours exposure to 10 g/mL of copper |
| SRR5124581 | + | + | PRJNA358429 | 2223 | olfactory tissue after 24 hours exposure to 10 g/mL of copper |
| SRR5124582 | + | + | PRJNA358429 | 3302 | olfactory tissue after 24 hours exposure to 0 g/mL of copper |
| SRR5124583 | + | + | PRJNA358429 | 2104 | olfactory tissue after 24 hours exposure to 10 g/mL of copper |
| SRR5312374 | + | + | PRJNA377918 | 17894 | spinal cord uninjured |
| SRR5312375 | + | + | PRJNA377918 | 2433 | spinal cord post injury 6 hours |
| SRR5312376 | + | + | PRJNA377918 | 5199 | spinal cord post injury 1 day |
| SRR5312378 | + | + | PRJNA377918 | 7179 | spinal cord post injury 1 week |
| SRR5312379 | + | + | PRJNA377918 | 10103 | spinal cord post injury 2 week |
| SRR5312380 | + | + | PRJNA377918 | 11737 | spinal cord post injury 3 week |
| SRR5312381 | + | + | PRJNA377918 | 8847 | spinal cord post injury 4 week |
| SRR5312382 | + | + | PRJNA377918 | 5257 | spinal cord post injury 5 week |
| SRR5312383 | + | + | PRJNA377918 | 6554 | spinal cord post injury 6 week |
| SRR5312384 | + | + | PRJNA377918 | 12586 | spinal cord post injury 12 week |
| SRR5312385 | + | + | PRJNA377918 | 14228 | brain uninjured |
| SRR5312386 | + | + | PRJNA377918 | 7693 | brain post injury 6 hours |
| SRR5312387 | + | + | PRJNA377918 | 6284 | brain post injury 1 day |
| SRR5312388 | + | + | PRJNA377918 | 9861 | brain post injury 3 days |
| SRR5312389 | + | + | PRJNA377918 | 7860 | brain post injury 1 week |

| | | | | | |
|---|---|---|---|---|---|
| SRR5312390 | + | + | PRJNA377918 | 7615 | brain post injury 2 week |
| SRR5312391 | + | + | PRJNA377918 | 9491 | brain post injury 3 week |
| SRR5312392 | + | + | PRJNA377918 | 10065 | brain post injury 4 week |
| SRR5312393 | + | + | PRJNA377918 | 6349 | brain post injury 5 week |
| SRR5312394 | + | + | PRJNA377918 | 6347 | brain post injury 6 week |
| SRR5312395 | + | + | PRJNA377918 | 10879 | brain post injury 12 week |
| SRR5431797 | + | - | PRJNA382290 | 564 | Juvenile liver |
| SRR5431798 | + | - | PRJNA382290 | 578 | liver at metamorphesis stage 5 |
| SRR5431799 | + | - | PRJNA382290 | 580 | liver at metamorphesis stage 2 |
| SRR5431800 | + | - | PRJNA382290 | 574 | larvae liver |

**Table S4**. Selection analyses on mammal ALK and LTK using different models of PAML. Kappa ($\kappa$) is the transition/transversion rate; $p_i$ is the proportion of sites inferred to be in each selection category $i$; df is degrees of freedom; ln $L$ is the natural log of the likelihood.

**Table S4.1**. Levels of selection ($\omega$) within each clade, calculated under clade model C.

| | | | Purifying selection class | | Neutral site class | | Divergent selection site class | | ln $L$ | $P$ value[a] (df) |
|---|---|---|---|---|---|---|---|---|---|---|
| Clade | Tree length | $k$ | $\omega_0$ | $p_0$ | $\omega_1$ | $p_1$ | $\omega_2$ $\omega_3$ $\omega_4$ | $p_2$ | | |
| ALK | 42.5 | 1.95 | 0.039 | 0.37 | 1 | 0.17 | 0.319 0.171 | 0.46 | -89260.6 | < 0.01 (1) |
| LTK | 43.8 | 1.96 | 0.28 | 0.48 | 1 | 0.15 | 0.029 0.109 | 0.38 | -89185.5 | < 0.01 (1) |
| ALK + LTK | 43.4 | 1.95 | 0.29 | 0.48 | 1 | 0.15 | 0.032 0.014 0.110 | 0.38 | -89169.6 | < 0.01 (2) |
| $M_{2a\_rel}$ | 44.7 | 1.96 | 0.28 | 0.47 | 1 | 0.161 | 0.037 | 0.37 | -89313.5 | |

[a] $M_{2a\_rel}$ was used as the null model for the likelihood ratio test, as in Weadick and Chang (2012). For the null model $M_{2a\_rel}$, no foreground was defined.

**Table S4.2**. Levels of selection ($\omega$) as determined by execution of branch models (PAML; 100).

| Foreground branches[a] | Foreground $\omega_f$ | Background $\omega_b$ | ln $L$ | $P$ value |
|---|---|---|---|---|
| — | 0.173 | | -92651.6 | |
| ALK | 0.152 | 0.181 | -92643.9 | < 0.01 |
| LTK | 0.259 | 0.162 | -92619.1 | < 0.01 |
| ALK + LTK | 0.153 0.258 | 0.167 | -92617.2 | < 0.01 |

[a] Nonsynonymous:synonymous substitution ratio $\omega$ was permitted to vary among branches in the phylogeny by designating the foreground branches and using the rest of the branches in the phylogeny as background. For the null model $M_0$ (first row), no foreground was defined.

**Table S4.3** Levels of selection ($\omega$) from models that allow the ratio $\omega$ to vary among sites.

| Clade | Model[a] | ln $L$ | Parameter estimates[b] | Models Compared | $P$ value[c] | Sites under selection[d] |
|---|---|---|---|---|---|---|
| ALK | $M_0$ | -92651.6 | $\omega_0 = 0.173$; tree length = 33.85; $\kappa = 1.79$ | | | |
| | $M_{1a}$ | -90488.6 | $p_0 = 0.557$; $\omega_0 = 0.11$ | $M_{1a}$:$M_{2a}$ | 1 | 901 |
| | $M_{2a}$ | -90488.6 | $p_0 = 0.56$; $\omega_0 = 0.11$; $p_2 = 0.071$; $\omega_2 = 1.00$ | | | |
| | $M_7$ | -89118.9 | $p = 0.61$; $q = 1.71$ | $M_7$:$M_8$ | < 0.01 | 901 |
| | $M_8$ | -89110.2 | $p_0 = 0.99$; ($p_1 = 0.009$); $\omega = 2.84$ | | | |
| LTK | $M_0$ | -92651.6 | $\omega_0 = 0.17$; tree length = 33.85; $\kappa = 1.79$ | | | |
| | $M_{1a}$ | -90488.6 | $p_0 = 0.557$; $\omega_0 = 0.11$ | $M_{1a}$:$M_{2a}$ | 1 | 901 |
| | $M_{2a}$ | -90488.6 | $p_0 = 0.557$; $\omega_0 = 0.11$; $p_2 = 0.13$; $\omega_2 = 1.00$ | | | |
| | $M_7$ | -89118.9 | $p = 0.61$; $q = 1.71$ | $M_7$:$M_8$ | < 0.01 | |
| | $M_8$ | -89110.2 | $p_0 = 0.99$; ($p_1 = 0.009$); $\omega = 2.84$ | | | |

[a] $M_0$: one-ratio model; $M_{1a}$: neutral model; $M_{2a}$: selection model; $M_7$: Beta; $M_8$: Beta & ω.
[b] Parameters $p$, $p_0$, $p_1$, $p_2$, and $q$ govern the shapes of beta distributions within models.
[c] Likelihood-ratio tests were performed to test whether $M_2$ and $M_8$ models fit the data better than $M_1$ and $M_7$ respectively.
[d] Site(s) under significant positive selection (Bayesian posterior probability > 0.95) , identified via Bayes Empirical Bayes analysis (104).

**Table S5**. Pairwise evolutionary rates estimated for ALK and LTK for representatives of mammals, fishes, and other vertebrates.

| Category | Species | ALK[a] | LTK[a] | Comparison of rates |
|---|---|---|---|---|
| Mammals | human vs opossum | 0.103 | 0.244 | ALK < LTK $P < 0.01$ |
| | vervet monkey vs opossum | 0.099 | 0.247 | |
| | armadillo vs opossum | 0.073 | 0.418 | |
| | armadillo vs human | 0.064 | 0.530 | |
| | armadillo vs vervet monkey | 0.063 | 0.533 | |
| | mouse vs opossum | 0.099 | 0.239 | |
| | mouse vs human | 0.073 | 0.273 | |
| | mouse vs vervet monkey | 0.066 | 0.272 | |
| | mouse vs armadillo | 0.057 | 0.450 | |
| Fish | zebrafish vs ghostshark | 0.120 | 0.172 | ALK > LTK $P = 0.14$ |
| | *Latimeria* vs ghostshark | 0.186 | 0.150 | |
| | Spotted gar vs ghostshark | 0.317 | 0.139 | |
| | Spotted gar vs *Latimeria* | 0.186 | 0.125 | |
| Others | *Anolis* vs zebrafinch | 0.173 | 0.163 | ALK = LTK $P = 0.47$ |
| | *Xenopus* vs zebrafinch | 0.145 | 0.155 | |
| | *Xenopus* vs *Anolis* | 0.146 | 0.172 | |

[a] dN/dS, calculated by the LPB93 method using `PAML` (114).
